# Exclusion of condensin I from the nucleus during prophase coordinates mitotic chromosome reorganization to complete sister chromatid resolution

**DOI:** 10.1101/2024.04.26.591320

**Authors:** John K. Eykelenboom, Marek Gierliński, Zuojun Yue, Tomoyuki U. Tanaka

## Abstract

During early mitosis, chromosomes are reorganized from their relatively decompacted interphase state into characteristic mitotic rod-shaped structures. This process is important to allow correct segregation of newly replicated sister chromatids to the opposite spindle poles during anaphase. To facilitate mitotic chromosome organization, two protein complexes named condensin I and condensin II play crucial roles. Condensin II is particularly important for achieving sister chromatid separation (resolution) whilst condensin I is required for chromosome condensation (compaction). Although sister chromatid resolution occurs 15-20 min earlier before chromosome compaction, it is not yet clear how these events are temporally coordinated or whether this temporal coordination is important to ensure chromosome segregation later in mitosis. One hypothesis is that the temporal coordination is achieved through different subcellular localisation of two condensin complexes; whilst condensin II localizes in the nucleus, condensin I is restricted to the cytoplasm, during interphase and prophase. In this study we tested this hypothesis by engineering the localization of condensin I to the nucleus. We monitored sister chromatid resolution and chromosome compaction by real-time imaging that visualized selected neighboring chromosome loci. We found that localization of condensin I to the nucleus led to precocious chromosome compaction during prophase with a similar timing to sister chromatid resolution. We also monitored later stages of mitosis and found that cells expressing nuclear condensin I subsequently exhibited frequent chromosome mis-segregation in anaphase. Therein, the majority of mis-segregated chromosomes consisted of lagging chromosomes involving both sister chromatids. This suggests that the temporal control of mitotic chromosome reorganization is crucial for high-fidelity chromosome segregation. In conclusion, the exclusion of condensin I from the nucleus during prophase delays chromosome compaction and allows condensin II to complete sister chromatid resolution, which ensures correct chromosome segregation later in mitosis.

## Introduction

Segregation of chromosomes in mitosis is preceded by the reorganisation of chromosomes into their characteristic mitotic shapes in human cells. This chromosome reorganization takes place during early mitosis (prophase and prometaphase) and involves two major structural changes: First, sister chromatids are resolved from each other while both sister chromatid cohesion and DNA catenation are removed (sister chromatid resolution). Second, each chromatid is compacted both axially and laterally (chromosome compaction). These structural changes are required for correct chromosome segregation during the subsequent anaphase (Spicer and Gerlich, 2023; Swedlow and Hirano, 2003; Takahashi and Hirota, 2019). In terms of timing, sister chromatid resolution happens 15-20 min earlier than chromosome compaction in human cells (Eykelenboom et al., 2019). However, it is still unknown how sister chromatid resolution and chromosome compaction are temporally coordinated or whether such coordination is important for correct chromosome segregation.

Mitotic chromosome reorganisation is dependent on protein complexes known as the condensin complexes (Hirano, 2016; Uhlmann, 2016). The condensin complexes belong to a family of <u>s</u>tructural <u>m</u>aintenance of <u>c</u>hromosome (SMC) complexes. Like other SMC- complexes, the condensin complex forms a tripartite ring structure, which consists of two SMC proteins (SMC2 and SMC4) and a Kleisin subunit. In addition, the condensin complex contains two kleisin-associating proteins termed Hawks (<u>H</u>EAT proteins <u>a</u>ssociated <u>w</u>ith <u>K</u>leisins) (Hirano, 2016; Uhlmann, 2016; Wells et al., 2017). The condensin complex promotes the capture and extrusion of a linear chromatin to form a chromatin loop. This process is dependent on the activity of two ATPase domains within the globular heads of SMC-proteins (Davidson et al., 2019; Ganji et al., 2018; Pradhan et al., 2023). It is suggested that such chromatin loop formation drives the mitotic chromosome reorganization (Gibcus et al., 2018; Naumova et al., 2013).

Although some eukaryotes have only a single condensin complex, the majority of multicellular organisms (including plants and most animals) contain two distinct condensin complexes termed condensin I and condensin II (Hoencamp et al., 2021). These two complexes both contain the same SMC2 and SMC4 components, but different Kleisin (NCAPH in condensin I; NCAPH2 in condensin II) and Hawk proteins (NCAPG and NCAPD2 in condensin I; NCAPG2 and NCAPD3 in condensin II) (Figure 1A) (Hirano et al., 1997; Hirano and Mitchison, 1994; Ono et al., 2003). Despite their overall structural similarity, loss of either the condensin I or II complex from the cell yields distinct phenotypes, suggesting they perform different tasks (Ono et al., 2003). For example, loss of condensin II leads to generation of stretched chromosomes lacking axial rigidity and severe chromosome segregation defects. In contrast, loss of condensin I leads to more subtle segregation defects but yields chromosomes which are wider and shorter and with a more disorganised central core (Green et al., 2012). More recently, it was suggested that condensin II is most important for untangling replicated sister chromatids early in mitosis (Eykelenboom et al., 2019; Nagasaka et al., 2016) and condensin I is more important during later stages of mitosis to achieve an organised compacted state (Eykelenboom et al., 2019; Gibcus et al., 2018). Both condensin I and II are activated in early mitosis through the phosphorylation of their Kleisin and Hawk components by mitotic kinases such as cyclinB–Cdk1, AuroraB and Mps1 (Abe et al., 2011; Kagami et al., 2014; Kimura et al., 2001; Kimura et al., 1998; Lipp et al., 2007; Shintomi et al., 2015). Furthermore, recent studies suggest a detailed molecular picture of mitotic chromosomes, where condensin II forms relatively large chromatin loops (spanning ∼400 kb) around a central core and condensin I forms a series of shorter nested loops (spanning ∼80 kb) within the large loops (Gibcus et al., 2018; Walther et al., 2018). It is thought that the size difference between condensin I- and condensin II-dependent chromatin loops leads to their differential roles in the mitotic chromosome re-organization (Batty and Gerlich, 2019).

**Figure 1:**
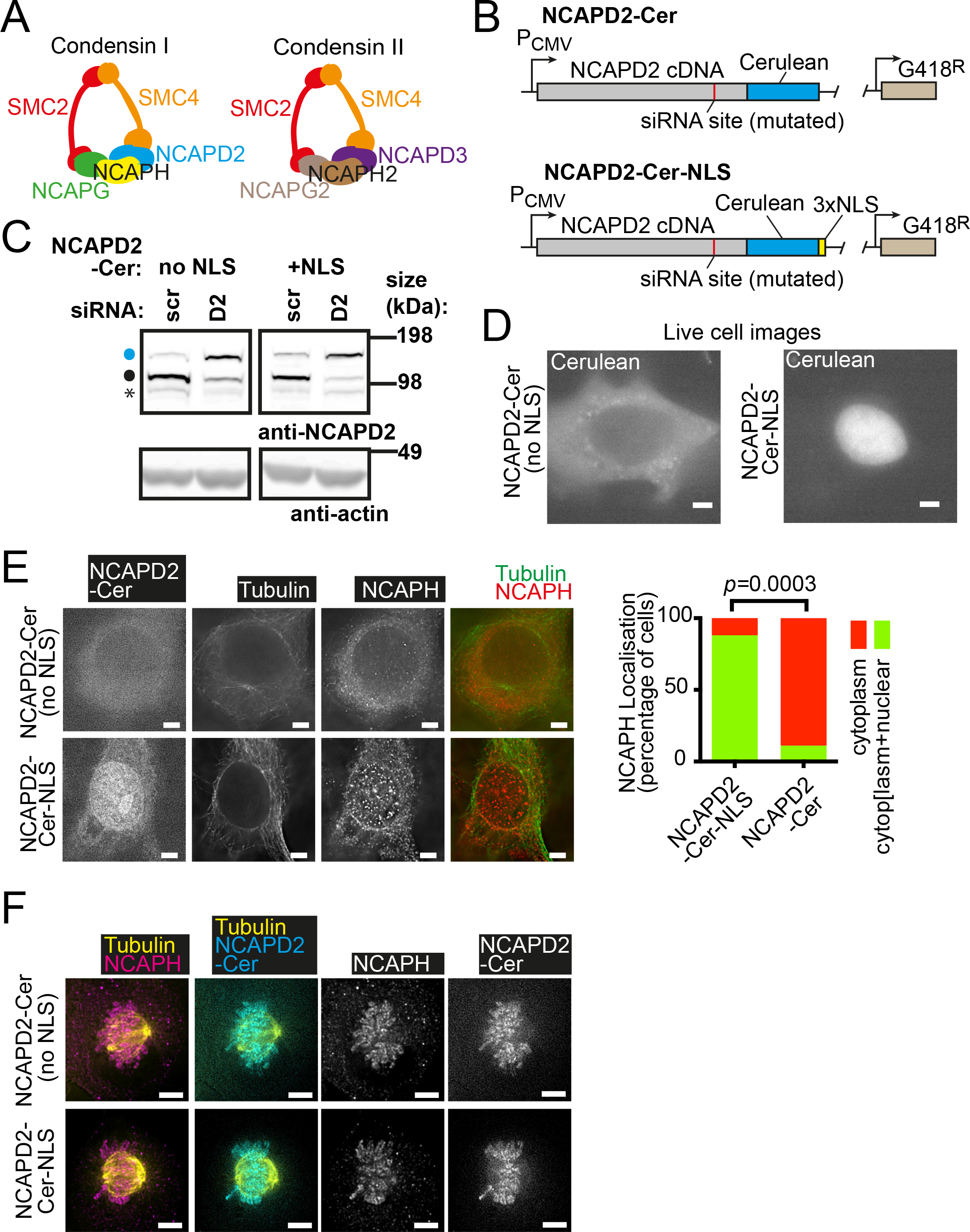
Generation of cell lines expressing Cerulean-tagged NCAPD2 (condensin I) in the cytoplasm or the nucleus of cells. (A) Cartoon depicting the proteins that make up condensin I and condensin II complexes. Note that SMC2 and SMC4 are common between the two whilst the other 3 subunits are unique to the specific complex. **(B)** Diagram depicting the strategy for expression of NCAPD2 that resides in the cytoplasm (top) or nucleus (bottom). The NCAPD2 cDNA was cloned under the control of a constitutive promoter (PCMV) in a plasmid containing the G418 resistance gene. Each construct was tagged at the C-terminus with the Cerulean fluorescent protein. One version of the construct had three copies of a nuclear localisation signal (NLS) at the very C-terminus (bottom) whilst the other did not (top). Moreover, the short siRNA target region was mutated to render these constructs insensitive to an siRNA against endogenous NCAPD2. **(C)** Western blot with an anti-NCAPD2 antibody for proteins from cells after treatment with NCAPD2 siRNA for 48 hours. The endogenous protein (NCAPD2; sensitive to the siRNA) is indicated by a small black circle whilst NCAPD2 tagged with Cerulean (NCAPD2-Cer; insensitive to the siRNA) is indicated by a blue circle. Actin is shown as a loading control. An asterisk indicates a non-specific band. **(D)** Live-cell visualisation of NCAPD2-Cerulean without or with the NLS sequence (left and right respectively) after transfection of HT-1080 cells with the plasmid constructs shown in part B. Scale bar 5 µm. **(E)** Localization of condensin I subunits in fixed interphase cells stably expressing either NCAPD2-Cer (top) or NCAPD2-Cer-NLS (bottom) after treatment with NCAPD2 siRNA for 48 hours. NCAPD2-Cerulean was visualised directly whilst NCAPH and Tubulin were visualised with immunofluorescence staining. The top and bottom panels show typical interphase cells displaying cytoplasmic or nuclear/cytoplasmic localisation of NCAPH, respectively. Scale bar 5 µm. The graph on the right-hand side shows quantification of these patterns for the indicated stable cell lines. The *p* value was obtained using a chi-square test. n = 14 for each condition. **(F)** Fixed metaphase cells expressing NCAPD2-Cer (top) or NCAPD2-Cer-NLS (bottom). NCAPD2-Cer and NCAPH were visualized as in E, on metaphase chromosomes. Scale bar 5µm.

Whilst condensin II and I promote sister chromatid resolution and chromosome compaction respectively, it is still unclear how the two complexes temporally coordinate these processes or whether such coordination is important for high-fidelity chromosome segregation later in mitosis. One possibility is that the differential subcellular localization of condensin I and II plays important roles: Whilst condensin II localizes in the nucleus throughout the cell cycle, condensin I resides in the cytoplasm during interphase and prophase, and only gains access to the chromosomes once the nucleus breaks down (Gerlich et al., 2006; Hirota et al., 2004; Ono et al., 2004). This means that condensin II, but not condensin I, has access to chromosomes during prophase when sister chromatid resolution begins. This may explain how sister chromatid resolution takes place 15-20 min earlier than chromosome compaction (Eykelenboom et al., 2019). In the present study, we formally test this possibility using a live- cell assay where we can analyze the kinetics of both sister chromatid resolution and chromosome compaction (Eykelenboom et al., 2019). We also address whether the temporal coordination of the two events is important for correct chromosome segregation during the subsequent anaphase.

## Results

### Engineered localization of condensin I to the nucleus

To address the functional importance of the exclusion of condensin I from the nucleus during interphase and prophase, we engineered the localization of condensin I to the nucleus. For this, we generated a stable HT-1080 diploid cell line which expressed a copy of the condensin I specific subunit NCAPD2 containing three copies of a nuclear localisation signal (NLS) at its C-terminus (Figure 1B, bottom). To visualize this protein in cells, we tagged it with the Cerulean fluorescent protein (NCAPD2-Cer-NLS). We also generated a control cell line expressing the same NCAPD2 construct but without NLSs (NCAPD2-Cer; Figure 1B, top). These constructs had silent mutations, which made them resistant to siRNA when siRNA was used to deplete the endogenous NCAPD2.

Intriguingly, when the endogenous NCAPD2 was depleted with siRNA, the expression levels of NCAPD2-Cer and NCAPD2-Cer-NLS were enhanced (Figure 1C, S1A). We reasoned that the reduced level of the endogenous protein allowed the incorporation of a higher level of NCAPD2-Cer and NCAPD2-Cer-NLS proteins into the condensin I complex, which stabilized these proteins. As expected, during interphase and prophase, NCAPD2-Cer-NLS localized predominantly in the nucleus while NCAPD2-Cer localized predominantly in the cytoplasm (Figure 1D). Importantly, when NCAPD2-Cer-NLS was expressed, other condensin I subunits NCAPH and NCAPG also showed a higher level of nuclear localization during interphase, suggesting that the whole condensin I complex changed its localization to the nucleus (Figure 1E, S1B, C). Moreover, in prometaphase/metaphase, both NCAPD2-Cer and NCAPD2-Cer-NLS (together with NCAPH and NCAPG) localized on condensed chromosomes (Figure 1F, S1D), suggesting that both proteins can form part of a functional condensin I complex.

### Exclusion of condensin I from the nucleus prevents precocious chromosome compaction during prophase

To analyse sister chromatid resolution and chromosome compaction, we previously developed a novel real-time assay in live cells (Eykelenboom et al., 2019). In this assay, *lac* and *tet*-operator arrays were integrated into two chromosome loci, 245 kb apart on human chromosome 5 (Figure 2A). These arrays were visualized as small GFP and mCherry fluorescent dots as they were bound by Lac repressor (with NLS) fused to GFP (GFP-LacI-NLS) and Tet repressor fused to mCherry, respectively. By observing the configuration of these dots over time, we were able to analyze how sister chromatid resolution and chromosome compaction proceed in early mitosis (Figure 2B, S2A). A unique advantage of this approach is that the kinetics of the two processes can be determined in a simple single assay. Normally, sister chromatid resolution occurs mainly during prophase (i.e. before nuclear envelope breakdown [NEBD]) and, after 15-20 min, chromosome compaction occurs during prometaphase (i.e. after NEBD) (Eykelenboom et al., 2019). However, when the level of NCAPD2 (a subunit of condensin I) was reduced, chromosome compaction was delayed (Eykelenboom et al 2019). On the other hand, when the level of NCAPD3 (a subunit of condensin II) was reduced, sister chromatid resolution was delayed.

**Figure 2:**
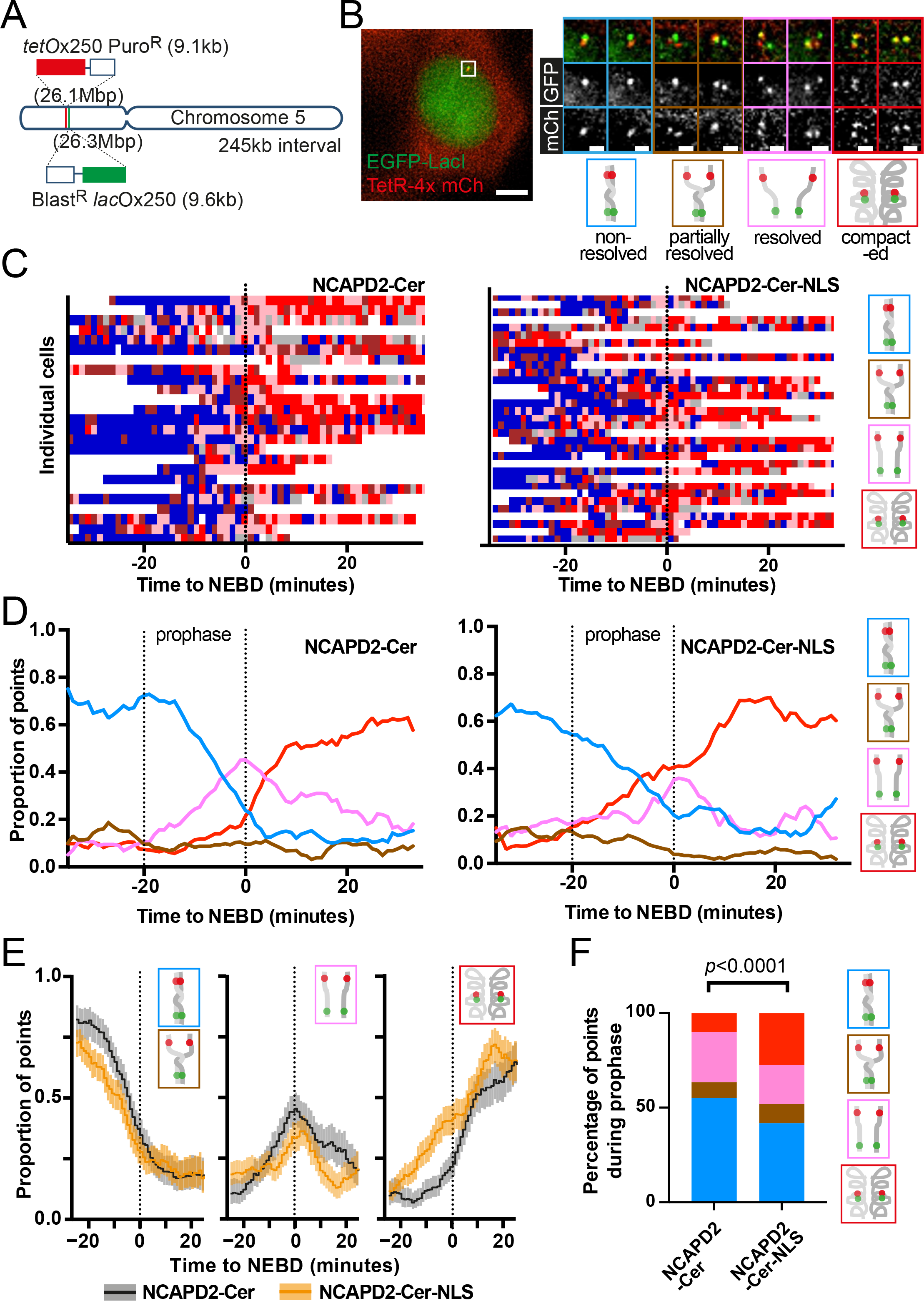
Access of condensin I to the nucleus during prophase leads to abnormal timing of mitotic chromosome organization (A) Diagram depicting the location of *tetO* and *lacO* arrays integrated into chromosome 5 (Eykelenboom et al., 2019). **(B)** Visualization of these arrays by expression of TetR-4x mCherry and EGFP-LacI in HT-1080 cells. The white box on the most left hand image indicates the operator arrays. The left-hand zoomed images are examples of the operator arrays adopting configurations/states characteristic of different stages during mitotic chromosome reorganisation (Eykelenboom et al., 2019); these states are indicated in the cartoons below. Designated color codes for each configuration are indicated in the frames of images and cartoons. Scale bars are 5µm (main cell) and 1µm (zoomed images). **(C)** Change in configuration of the operator arrays over time (*x* axis) as observed in individual live cells (each horizontal column represents each cell). Stable cell lines expressing either NCAPD2-Cer (left) or NCAPD2-Cer-NLS (right) were treated with an siRNA against endogenous NCAPD2 for 48 hours and synchronized by a double-thymidine block and release. Images were taken every minute, and, at each time point, the configuration of the operator arrays was determined as in B (color-coded as in B). Data from individual cells were aligned relative to NEBD (defined as time zero). **(D)** The proportion of each configuration (color-coded as in B) was determined from the data in C and plotted over time with smoothing (across 9 min). **(E)** The proportion of the blue or brown “not fully resolved” state, the pink “resolved” state and the red “compacted” state, observed before and after NEBD (-25 to 25 mins), for either NCAPD2-Cer cells (black line) or NCAPD2-Cer- NLS cells (orange line). The data was taken from C and D. The shaded areas of the plots indicate 95% confidence intervals determined using Clopper-Pearson method using a smoothing window of 10 minutes. **(F)** Graph shows the proportional representation of each state during prophase (-20 to 0 mins) for NCAPD2-Cer or NCAPD2-Cer-NLS cells. The data was taken from C and D. The *p* value was obtained using a chi-square test. n = 419 and 515 for NCAPD2-Cer and NCAPD2-Cer-NLS cells respectively. A more comprehensive analyses including different phases (G2 to late-prometaphase) is shown in Figure S2B.

We used this assay to evaluate the effect of engineered localization of condensin I to the nucleus on the timing of sister chromatid resolution and chromosome compaction. We took the cell lines with the GFP and mCherry fluorescent dots (Figure 2A, B), expressing NCAPD2-Cer and NCAPD2-Cer-NLS (Figure 1B) and depleted endogenous NCAPD2 by using a specific siRNA. In parallel, we synchronised these cells with a double thymidine block and a subsequent release. Approximately 8 to 10 hours after release from the second thymidine-induced arrest, we imaged the cells by microscopy to visualise the configuration of the GFP and mCherry dots over time. Cells that underwent NEBD were identified when signals of GFP-LacI-NLS (unbound to the *lac* operator array) became diffuse throughout the cell. In these cells, we identified the 3-dimensional coordinates of the GFP and mCherry dots and, based on this, we tracked how dot configuration changed over time relative to NEBD (time zero) in individual cells (Figure 2C). We then plotted the proportion of cells displaying each configuration over time (Figure 2D).

During late G2 phase (before -20 min, relative to NEBD), the GFP and mCherry dots often showed interchanges between non-resolved (2 dots; one of each colour; blue pattern) and partially resolved (3 dots; brown pattern) configurations, in both cell lines expressing either NCAPD2-Cer or NCAPD2-Cer-NLS (Figure 2C). Similar interchanges were observed during late G2 phase in wild-type cells expressing endogenous NCAPD2 only (without NCAPD2- Cer construct) (Eykelenboom et al., 2019). Then, during prophase (-20 to 0 min relative to NEBD) and prometaphase (after NEBD), more cells showed the resolved configuration (4 dots; two of each colour; pink pattern) and the compacted configuration (4 dots; 2 pairs of colocalised red and green dots) in both cell lines (Figure 2C). The cells expressing NCAPD2-Cer showed an increase in the resolved configuration (pink line) during prophase, followed by an increase in the compacted configuration (red line) mainly after NEBD (Figure 2D, left). This sequence of events was similar to that of wild-type cells (Eykelenboom et al., 2019). On the other hand, in the cells expressing NCAPD2-Cer-NLS, there was already an increase in the compacted configuration (red line) during prophase (Figure 2D, right). When the proportion of each configuration was compared between the cells with NCAPD2-Cer and NCAPD2-Cer-NLS (Figure 2E), a significant advancement of the compacted configuration was noticeable in cells with NCAPD2-Cer-NLS (Figure 2E, right). Consistent with this, the percentage of the compacted configuration (red) was higher during prophase and early prometaphase with NCAPD2-Cer-NLS than with NCAPD2-Cer (Figure 2F, S2B) though not at late G2 phase or late prometaphase (Figure S2B).

The above data suggest that, if condensin I localizes in the nucleus, it enhances chromosome compaction during prophase. Given that NCAPD2-Cer-NLS localizes in the nucleus during interphase and prophase, we next addressed if it also promotes chromosome condensation (compaction) prior to the entry into mitosis. To assess this, we measured the distance between the GFP and mCherry dots on the same chromatid – if two GFP dots and two mCherry dots were present, the two shorter GFP-mCherry dots’ distances were considered to be part of the same chromatids (Figure 3A). Intriguingly, the distance between the GFP and mCherry dots on the same chromatid was not significantly different between cells with NCAPD2-Cer or NCAPD2-Cer-NLS in late-G2 phase. However, this distance became significantly shorter during prophase and early prometaphase with NCAPD2-Cer- NLS (Figure 3B, C). The results suggest that, although condensin I was engineered to localize to the nucleus during interphase and prophase, it promoted chromosome compaction only during prophase, but not during G2 phase. This result is explained by previous reports that the activation of condensin I requires its phosphorylation by cyclin B- Cdk1 (Kimura et al., 2001; Kimura et al., 1998; Shintomi et al., 2015), which itself becomes activated and rapidly accumulates in the nucleus at prophase (Gavet and Pines, 2010a; Gavet and Pines, 2010b). Overall, we conclude that, in the context of wild-type cells, the exclusion of condensin I from the nucleus prevents premature chromosome compaction during prophase.

**Figure 3:**
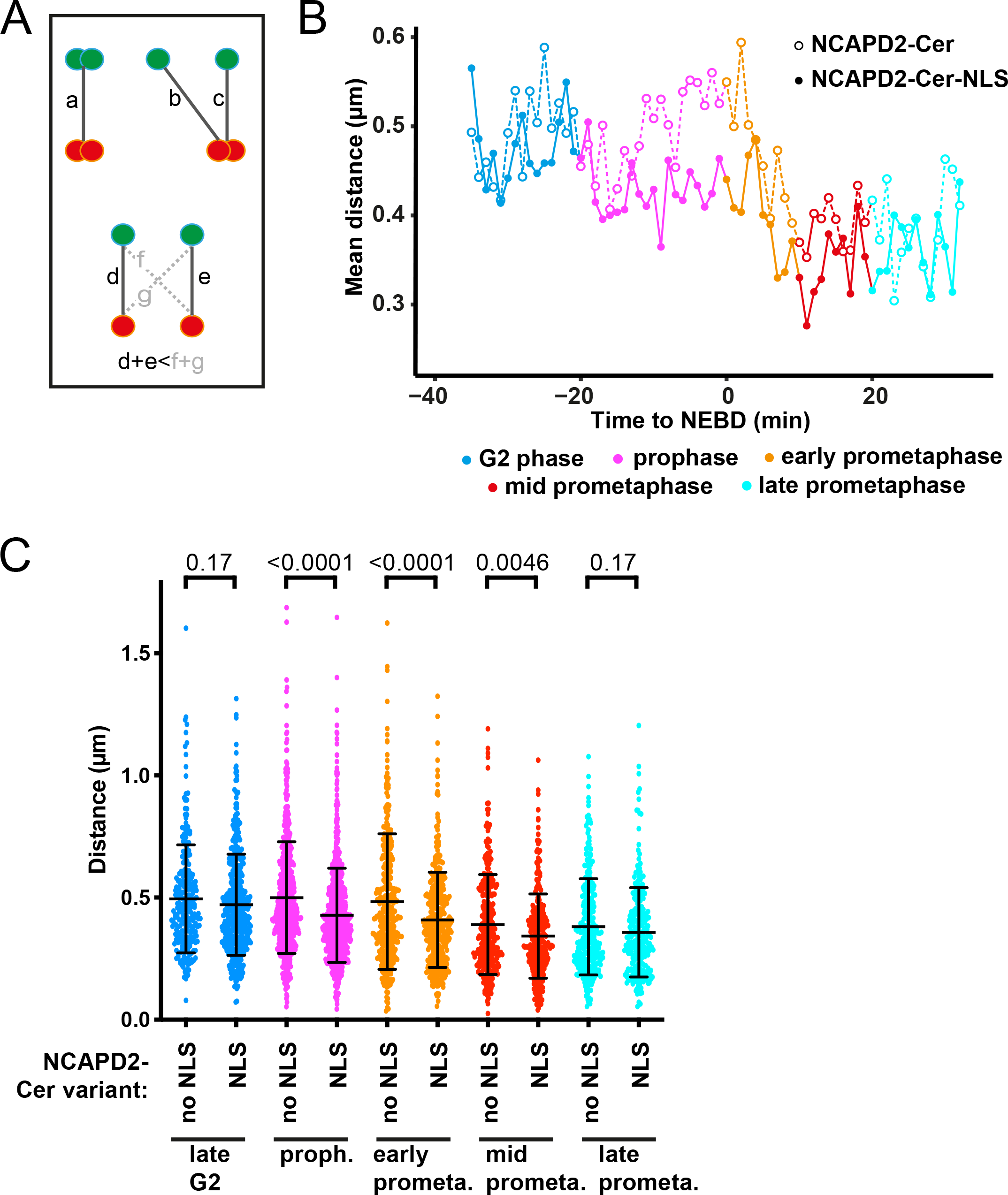
Access of condensin I to the nucleus during prophase leads to premature chromosome compaction in early stages of mitosis but not in G2 phase. (A) Diagram showing how chromosome compaction was measured with different configurations of operator arrays. Measurements were included from “non-resolved” (a), “partially resolved” (b and c) and “resolved/compacted” (d and e) configurations. Measurements f and g were excluded on the basis that *tetO* (red dot) and *lacO* (green dot) on the same chromatid are more likely found closer together. **(B)** Graph showing the mean distance between *tetO* and *lacO* (those indicated in A) in NCAPD2-Cer or NCAPD2-Cer-NLS cells over time during G2 and different stages of mitosis. The data are aligned relative to NEBD (defined as time zero). The measurements were obtained from cells imaged in Figure 1 C and D. Color coding of different phases is indicated below. **(C)** Graph showing individual *tetO-lacO* distances observed in NCAPD2-Cer or NCAPD2-Cer-NLS cells (also used in B) grouped for different G2 and mitotic phases (color-coded as in B). The mean and SD are shown for each data set. *p* values were obtained by comparing data between NCAPD2-Cer or NCAPD2-Cer-NLS cells at each phase using unpaired *t* tests. The number of analyzed data points for each data set was 252, 434, 575, 777, 360, 377, 285, 284, 270, 265 from left to right,

### Exclusion of condensin I from the nucleus during prophase subsequently ensures timely progression from metaphase to anaphase

We next addressed the effect of the engineered nuclear localization of condensin I on progression through mitosis. To do this, we used the HT-1080 cell lines we had already generated, which express LacI-GFP-NLS as well as either NCAPD2-Cer or NCAPD2-Cer- NLS. We depleted endogenous NCAPD2 by siRNA treatment and synchronised these cells using a double thymidine block. When we released the cells from this arrest, we added SiRDNA to the cell culture for fluorescent labelling of mitotic chromosomes. Approximately 10 hours after the release, we started imaging cells and observed them for 2.5 hours. We then focused on the cells that underwent NEBD during the observation (judged by diffusion of LacI-GFP-NLS from the nucleus to the whole cell) (Figure 4A). For each cell we measured the timings, relative to NEBD, of key mitotic events such as chromosome alignment on the metaphase plate (completion of chromosome congression), the onset of anaphase (start of chromosome segregation), and reformation of the nuclear envelope after chromosome segregation (judged by the re-accumulation of LacI-GFP-NLS into the nucleus) (Figure 4A, B). The progression from NEBD to chromosome alignment was similar between NCAPD2- Cer and NCAPD2-Cer-NLS cells. However, the anaphase onset (i.e. progression from metaphase to anaphase) was delayed in 14.8 % and 39.4 % with NCAPD2-Cer and NCAPD2-Cer-NLS, respectively (Figure 4B, C). In the control with scramble siRNA (without NCAPD2-Cer or NCAPD2-Cer-NLS), 17.0 % of cells showed a delay in the anaphase onset. Thus, the engineered nuclear localization of condensin I caused a delay in anaphase onset in some cells. In the context of wild-type cells, we conclude that the exclusion of condensin I from the nucleus during prophase subsequently ensures timely progression from metaphase to anaphase in a subset of cells.

**Figure 4:**
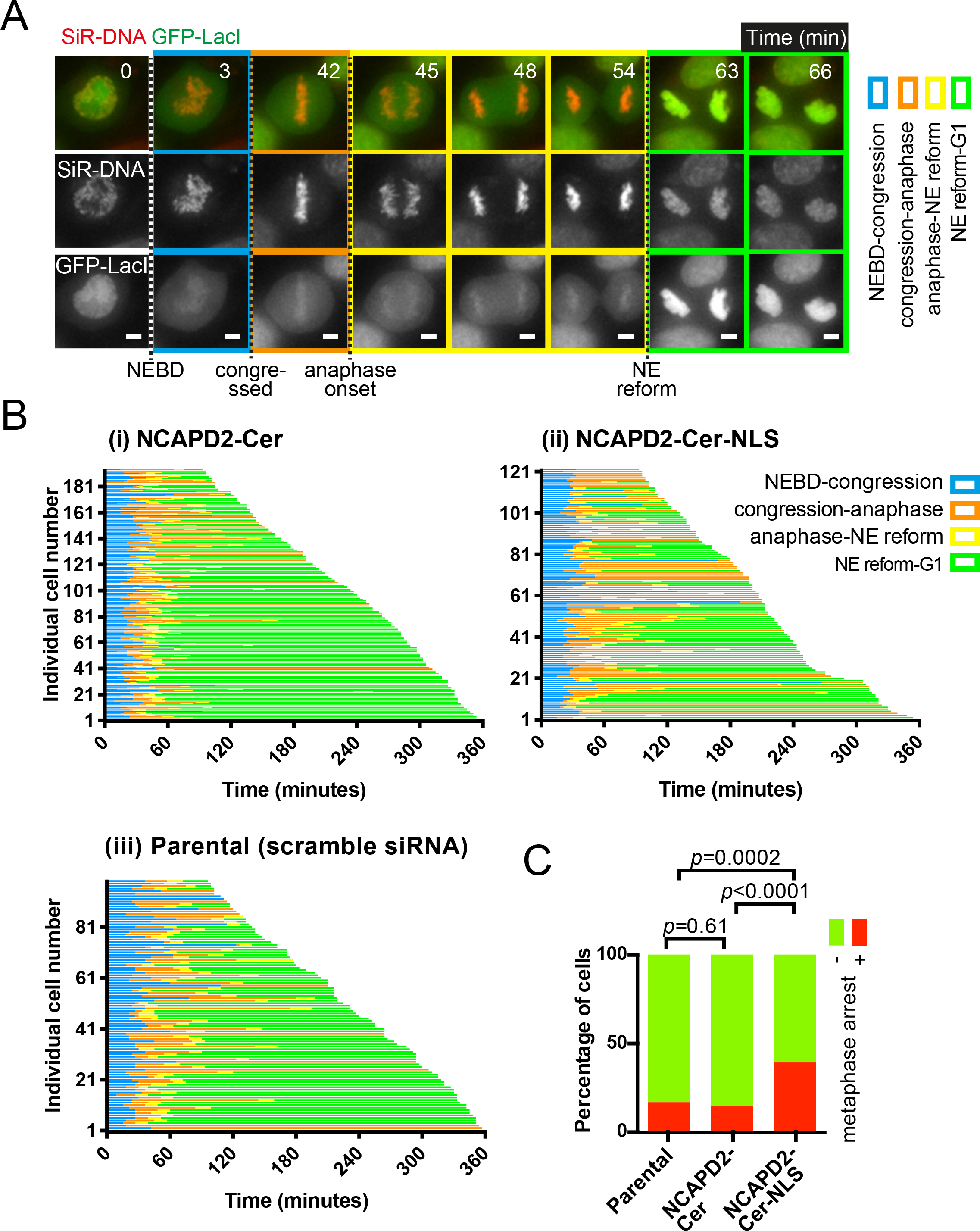
Disrupted mitotic chromosome reorganization results in delayed mitotic progression. (A) Time-lapse images show a representative cell passing through mitosis. Stable cell lines expressing either NCAPD2-Cer or NCAPD2-Cer-NLS were treated with an siRNA against endogenous NCAPD2 for 48 hours and synchronized by a double-thymidine block and released, approximately 10 hours after release images were taken every minute, and, different stages of mitosis were determined. NEBD was recognised by diffusion of GFP-LacI-NLS through the whole cell; congression by the alignment of all chromosomes (stained with SiRDNA) on the metaphase plate; anaphase by the first moment chromatids began segregating to opposite cell poles; and nuclear envelope reformation by relocalization of GFP-LacI-NLS back into the newly divided nuclei. Designated color codes for each stage are indicated in the image frames. Scale bars 5µm. The cell in this figure expressed NCAPD2-Cer, but mitotic progression of NCAPD2-Cer-NLS cells was also determined in the same way. **(B)** Mitotic progress of individual NCAPD2-Cer (i), NCAPD2-Cer-NLS (ii) or parental (iii) cells (*y* axis) plotted against time (*x* axis) and aligned relative to NEBD (defined as time zero). Cells expressing NCAPD2-Cer and NCAPD2-Cer-NLS were treated with siRNA against endogenous NCAPD2 whilst parental cells were treated with scramble siRNA. The colored lines, that use the same color codes as A, refer to trajectories of individual cells through mitosis. The end of the line indicates the end of the time-lapse for the cell. The number of analyzed cells for NCAPD2-Cer, NCAPD2-Cer-NLS and parental cells were 196, 127 and 100 respectively. **(C)** Quantification of the number of NCAPD2-Cer, NCAPD2-Cer-NLS or parental cells that did not progress further than metaphase during the time-lapse acquisition. The cells analysed were from B. The *p* values were obtained using Fisher’s exact test.

The transition from metaphase to anaphase in normal cells is subject to regulation by the spindle assembly checkpoint (SAC) (Lara-Gonzalez et al., 2021; McAinsh and Kops, 2023). We next tested if the delayed mitotic progression we had observed with NCAPD2-Cer-NLS was dependent on the SAC. We carried out a similar experiment to the previous one, except we included a low level of CPD-5, an inhibitor of the SAC regulator Mps1 kinase (Klaasen et al., 2022; Koch et al., 2016). The low concentration of CPD-5 was determined as the lowest concentration where anaphase progression was advanced but with minimal chromosome segregation defects (Figure S3A, B). With CPD-5, we found that almost all NCAPD2-Cer cells and NCAPD2-Cer-NLS cells now progressed to anaphase without difference between the two (Figure S3B-D). This suggests that NCAPD2-NLS led to a SAC-dependent delay in the anaphase onset in a subset of cells. Nonetheless, the majority (60.6%) of cells with NCAPD2-Cer-NLS still underwent timely entry into anaphase (Figure 4B, C).

### Exclusion of condensin I from the nucleus during prophase ensures correct chromosome segregation during the subsequent anaphase

We next studied the process of chromosome segregation in the cells with NCAPD2-Cer and NCAPD2-Cer-NLS, focusing on those that had shown normal timing in the metaphase-to-anaphase transition. Before observation, we depleted endogenous NCAPD2 by siRNA and then acquired images of these cells. In these experiments, we also included the parental cell line, i.e. without NCAPD2-Cer or NCAPD2-Cer-NLS and without depletion of endogenous NCAPD2. As shown in Figure 5, we observed cells undergoing anaphase and classified them into three different categories; (1) those that displayed normal chromosome segregation and subsequently formed two new daughter nuclei (Figure 5A); (2) those that showed lagging chromosomes, frequently leading to generation of micronuclei when the nuclear envelope reformed (Figure 5B, orange frame; inset ii, micronucleus); (3) those that showed defective nuclear segregation with several lagging or bridged chromosomes, followed by nuclear envelope reformation before nuclear segregation was completed (Figure 5B, red frame). We found that, while 16% of the parental or NCAPD2-Cer control cells showed defects in chromosome segregation (categories 2 and 3 above), 38% of NCAPD2- Cer-NLS cells showed such defects (Figure 5C). Moreover, if we consider category 3 alone, i.e. defects in the nuclear segregation, the difference between the parental/NCAPD2-Cer and NCAPD2-Cer-NLS was greater (3% vs 12%, respectively). To check reproducibility of the result, we repeated this experiment using alternative cell lines expressing NCAPD2-Cer and NCAPD2-Cer-NLS – we obtained similar results to Figure 5 (Figure S4A). In the context of wild-type cells, we conclude that the exclusion of condensin I from the nucleus during prophase ensures correct chromosome segregation during the subsequent anaphase.

**Figure 5:**
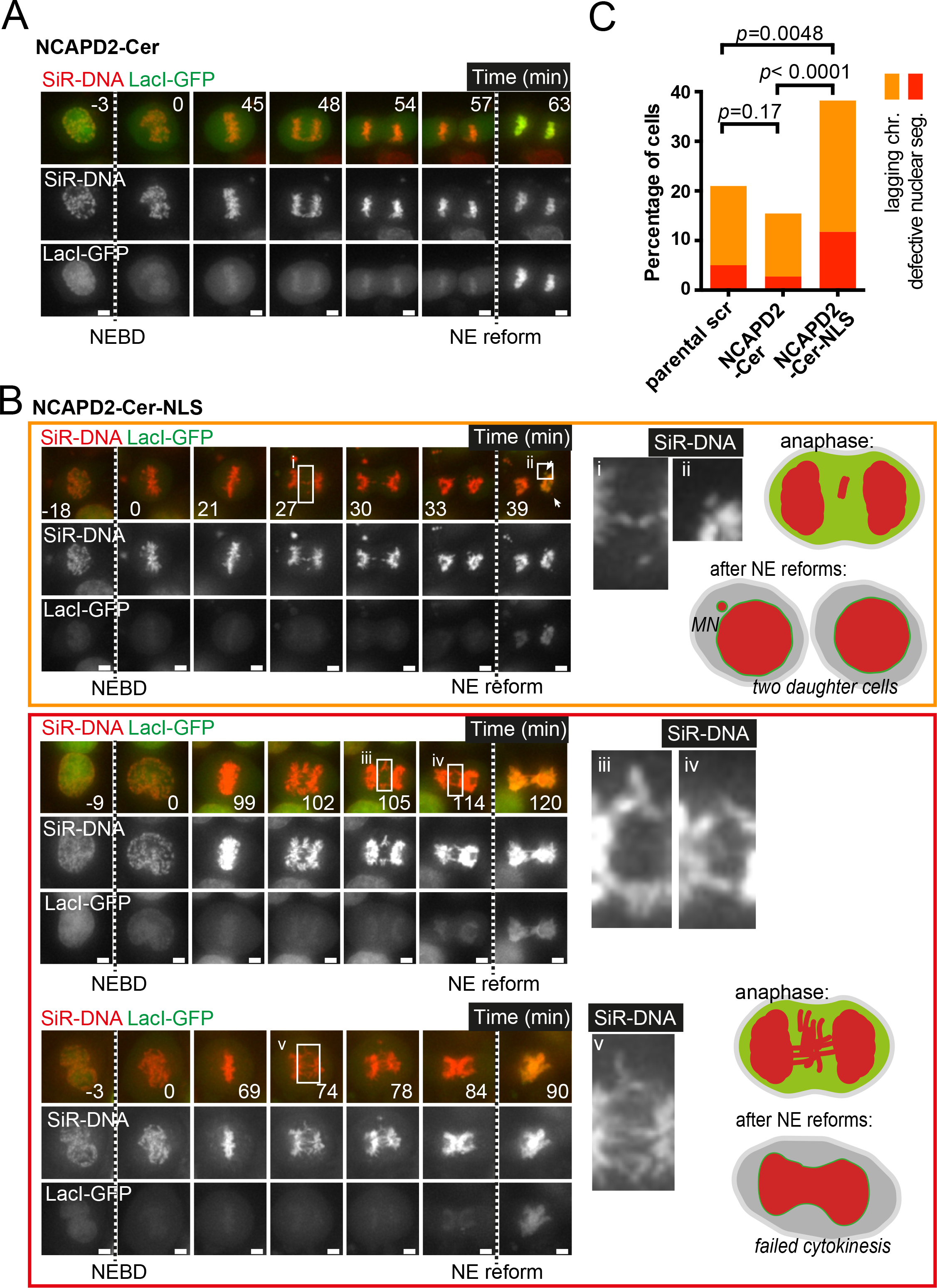
Disrupted mitotic chromosome reorganization results in chromosome segregation errors during anaphase. (A) Time-lapse images show a representative NCAPD2-Cer cell passing through mitosis. Cells were treated with an siRNA against endogenous NCAPD2 for 48 hours and synchronized by a double-thymidine block and released, approximately 10 hours after release images were taken every minute, and anaphase was observed. NEBD and the nuclear envelope (NE) reform were identified as in Figure 4A. Chromosomes were visualized using SiRDNA. **(B)** Sequences of time-lapse images show three different NCAPD2-Cer-NLS cells with segregation defects during anaphase. Cells were treated and imaged, and stages of mitosis were determined as in A. The upper panel, framed in orange, highlights a cell displaying lagging chromosomes in anaphase (see inset i) that later form a micronucleus (MN) after NE reformation (see inset ii). The lower panel, framed in red, highlights two cells with segregation defects that involve multiple sets of lagging chromosomes and lead to defective nuclear segregation (see insets iii, iv and v). These defects often culminate in the NE reform before complete nuclear segregation, leading to failure in cytokinesis. The cartoons to the right-hand side and within the colored frames summarize the observed defects. **(C)** Quantification of chromosome segregation defects for NCAPD2-Cer, NCAPD2-Cer-NLS or parental cells. Cells were treated, imaged and categorized according to A and B. Parental cells were treated with scramble siRNA. In each cell line, the cells that performed anaphase without problem make up the remainder percentages but are not shown on the graph. The *p* values were obtained using chi-square test for trends. The number of analyzed cells for NCAPD2-Cer, NCAPD2- Cer-NLS and parental cells were 181, 102 and 119 respectively.

In general, lagging chromosomes could be generated by various mechanisms; for example, merotelic attachments (where a single kinetochore attaches to microtubules from opposite spindle poles) would generate lagging chromosomes involving single chromatids (Gregan et al., 2011), while defects in either sister chromatid resolution or removal of sister chromatid cohesion would produce lagging chromosomes containing both sister chromatids (Batty and Gerlich, 2019; Shintomi and Hirano, 2010). When lagging chromosomes were generated with NCAPD2-Cer-NLS, it seemed that pairs of chromosomes, probably sister chromatids, were frequently involved (Figure 5B, inset i, iii and iv).

### Evidence that exclusion of condensin I from the nucleus during prophase ensures complete sister chromatid resolution

To characterize these lagging chromosomes further, we visualized the chromosomal DNA, the microtubules and the centromere marker ACA, by immuno-staining in fixed anaphase cells expressing NCAPD2-Cer or NCAPD2-Cer-NLS (where the endogenous NCAPD2 had been depleted by siRNA) (Figure 6A). As a control we also analyzed parental cells (without NCAPD2-Cer or NCAPD2-Cer-NLS and with no depletion of endogenous NCAPD2), treated with Reversine, an inhibitor of Aurora B and Mps1 kinases (Santaguida et al., 2010) that are both important for eliminating merotelic attachments (Cimini, 2007; Petsalaki and Zachos, 2013). Consistent with Figure 5C, while the NCAPD2-Cer cells showed a similar number of lagging chromosomes (ACA signals) to the parental cells, the NCAPD2-Cer-NLS cells showed a larger number of lagging chromosomes than the NCAPD2-Cer cells (Figure 6B). Moreover, the parental cells showed a larger number of lagging chromosomes (ACA signals) when treated with Reversine than without this treatment (Figure 6B). In these cells, when the number of lagging chromosomes was limited and they were spatially separated from each other, we were able to characterize them individually (Figure 6A, orange frame). By contrast, when their number was large, they became too crowded to distinguish them individually (Figure 6A, red frame). The NCAPD2-Cer-NLS cells showed a larger number of both discernible and indiscernible lagging chromosomes than did the NCAPD2-Cer cells (Figure 6C). Similarly, the parental cells showed a larger number of both discernible and indiscernible lagging chromosomes when treated with Reversine compared to when they were untreated (Figure 6C).

**Figure 6:**
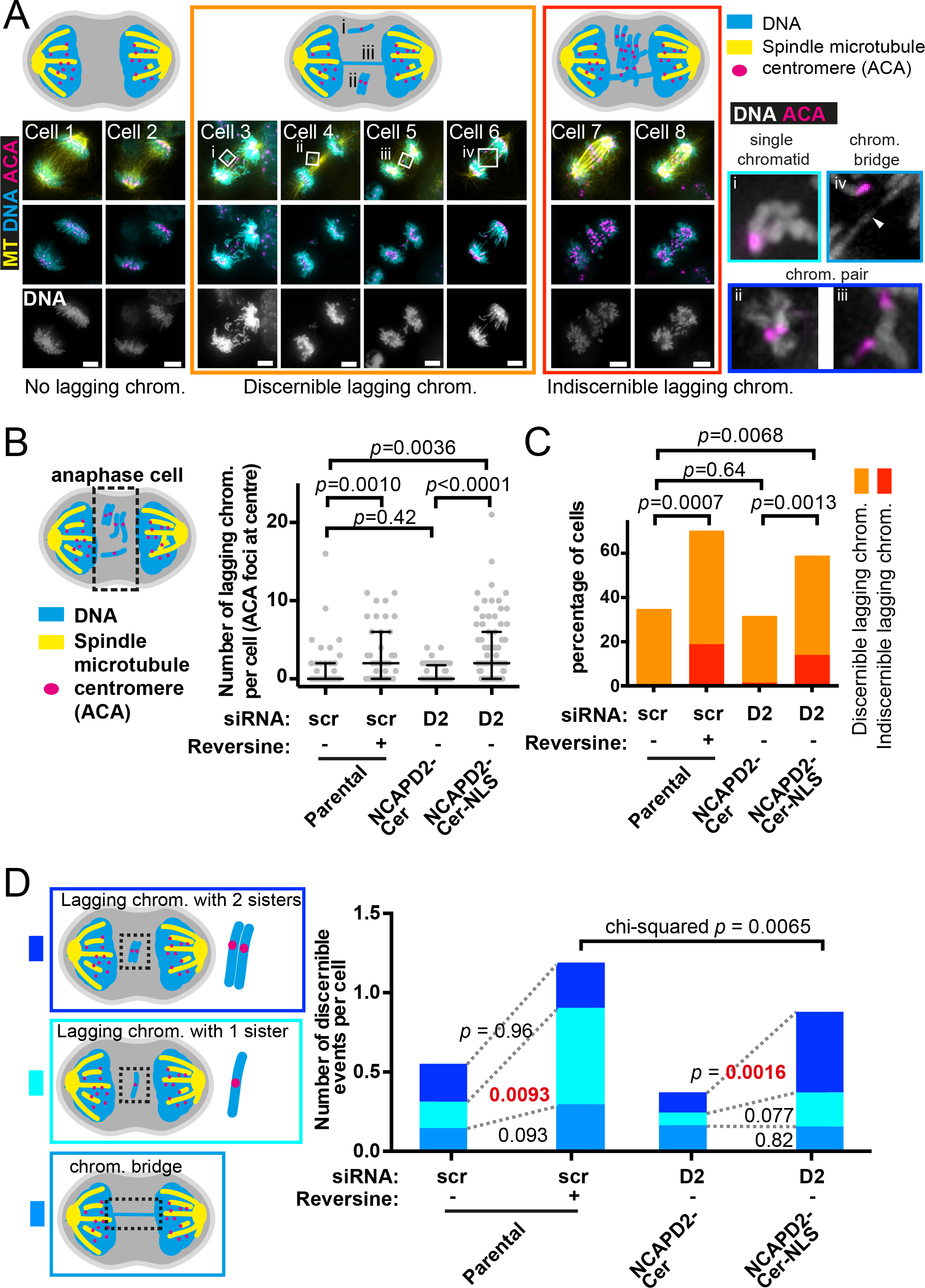
Mis-segregating chromosomes associated with nuclear localization of condensin I often contain two sister centromeres. (A) Fixed-cell analysis of centromeres and chromosomes during anaphase. NCAPD2-Cer or NCAPD2-Cer-NLS cells were treated with an siRNA against endogenous NCAPD2 for 48 hours and synchronized by a double- thymidine block and release. Approximately 10 hours after release, cells were fixed with paraformaldehyde and centromeres (ACA) and microtubules (MT) were stained by immunofluorescence and DNA was stained with DAPI. Anaphase cells were identified by the elongated spindle and segregating masses of DNA. Cells 1 and 2 showed normal anaphase, with all ACA signals localised with the two segregating masses of DNA. Cells 3 to 6 displayed individual isolated lagging chromosome(s) made up of (i) a single chromatid, identified by a single ACA foci on a DAPI stained object, (ii) and (iii) chromosome pairs, identified by two ACA foci on the same DAPI stained object or (iv) a chromosome bridge, identified as a DAPI stained object (indicated by a white arrowhead), stretched between the main segregating masses of DNA. Cells 7 and 8 displayed 6 or more lagging chromosomes with large DAPI stained objects including several ACA foci (masses of chromosomes) that lie between two segregating DNA masses. The cartoons above the images summarize these chromosome segregation defects. The orange and red colored frames surrounding the individual cell examples respectively indicate those in which lagging chromosome events can be individually identified (relatively low number of events) or not (relatively high number of events). The colored frames surrounding the zoomed insets to the right-hand side are color coded as in diagrams in D. Scale bars 5µm. **(B)** Quantification of the number of ACA foci found at the centre of the cell, between segregating chromosome masses (indicated in the cartoon to left-hand side), in parental cells treated or not with Reversine, or in NCAPD2- Cer or NCAPD2-Cer-NLS cells. Cells were treated as in A and, where applicable, Reversine was added for two hours before fixation. The median and interquartile range are shown for each cell line. The number of analyzed cells for NCAPD2-Cer or NCAPD2-Cer-NLS were 56 and 78 respectively. The number of analyzed untreated or Reversine treated parental cells were 43 and 37 respectively. The *p* values were obtained using Mann-Whitney U test. **(C)** Quantification of the percentage of cells exhibiting chromosome segregation defects, which are individually discernible (orange frame in A) or indiscernible (red frame in A). The data was taken from B. In each condition, the cells that displayed no problems make up the remainder percentages but are not shown on the graph. The *p* values were obtained using chi-square test. **(D)** Characterizing discernible lagging chromosomes (those indicated by the orange frame in A). The data was taken from images of cells acquired in B. The observed discernible lagging chromosomes were classified as in the cartoon on the left-hand side and as exemplified in A (right-hand side zoomed images). The number of events per cell was derived by dividing the number of classified events by the number of cells that were included in this analysis. Because cells with several indiscernible events were not included in this analysis, the values do not reflect the actual number of lagging chromosome per cell. Statistical tests and *p* values were obtained with Mann-Whitney U tests unless indicated otherwise. The number of analyzed cells/number of events for NCAPD2-Cer or NCAPD2- Cer-NLS were 63/23 and 65/57 respectively. The number of analyzed cells/number of events for untreated or Reversine treated parental cells were 42/24 and 28/33 respectively.

When we were able to discern lagging chromosomes individually (Figure 6A, orange frame), we found that there were three categories of lagging chromosomes; 1) a lagging chromosome consisting of a single chromatid (with a single ACA signal) (Figure 6A, inset i), 2) lagging chromosomes consisting of two sister chromatids (paired chromosomes with two ACA signals) (Figure 6A, inset ii, iii), and 3) a chromosome bridge (DAPI signal stretched between the two segregating masses of chromosomes, ACA signals with segregating centromeres) (Figure 6A, inset iv). The Reversine treatment of the parental cells mainly increased the number of lagging chromosomes consisting of single chromatids (Figure 6D), which is consistent with an increase in merotelic attachment (Gregan et al., 2011). By contrast, the NCAPD2-Cer-NLS cells showed a significantly higher number of lagging chromosomes involving both sister chromatids than did the NCAPD2-Cer cells (Figure 6D).

Lagging chromosomes involving sister chromatids can be generated by either incomplete sister chromatid resolution during early mitosis or through failure to establish or loss of microtubule attachments at both sister kinetochores. The latter possibility is unlikely because we detected microtubule attachments to both sister centromeres (ACA) in most cases of lagging sister chromatids, and no microtubule attachments to sister centromeres only in rare cases (Figure S4B). Therefore, we reason that with the engineered nuclear localization of condensin I, sister chromatid resolution was not completed in some chromosomes, causing more frequent chromosome missegregation. Thus, in the context of wild-type cells, we suggest that the exclusion of condensin I from the nucleus during prophase allows condensin II to complete sister chromatid resolution in all chromosomes.

## Discussion

### Sister chromatid resolution and chromosome compaction are temporally coordinated by subcellular condensin I localization to ensure correct chromosome segregation

The importance of condensin I and condensin II in mitotic chromosome organisation is long established through several studies. Condensin II plays an important role in sister chromatid resolution and condensin I in chromosome compaction (Eykelenboom et al., 2019; Gibcus et al., 2018; Nagasaka et al., 2016). Whilst a number of studies suggest that resolution precedes compaction it has still been unknown how this sequential timing is achieved (Eykelenboom et al., 2019; Nagasaka et al., 2016). One possibility is that this timing is defined by the exclusion of condensin I from the nucleus during the early stages of mitosis. In this study, we have directly addressed this possibility by observing both sister chromatid resolution and chromosome compaction in live cells expressing a version of condensin I that is present in the nucleus before NEBD. We found that with the engineered localization of condensin I in the nucleus, there is advancement of condensin I-mediated compaction of chromosomes in prophase such that it occurs almost simultaneously with sister chromatid resolution. Although in our system condensin I was also present in the nucleus during interphase, we did not observe precocious chromosome compaction in interphase, presumably because condensins are activated by CDK1 and other mitotic kinases only when cells reach prophase (Abe et al., 2011; Kimura et al., 2001; Kimura et al., 1998; Shintomi et al., 2015).

Our results also showed that precocious nuclear localization of condensin I led to frequent chromosome missegregation in anaphase, involving non-disjunction of sister chromatids. Although sister chromatid resolution still largely occurs in this condition, the completion of the resolution often seemed to be defective. This suggests that temporal coordination of sister chromatid resolution and chromosome compaction is crucial to ensure the completion of sister chromatid resolution for all chromosomes. Taken together we propose that condensin I has evolved to stay outside the nucleus to allow thorough sister chromosome resolution before compaction is instigated following NEBD.

### How the access of condensin I to chromosomes in prophase could interfere with completion of sister chromatid resolution

Although both condensin I and II complexes form loops from linear DNA molecules during mitosis (Kong et al., 2020), condensin II is known to form larger DNA loops than condensin I as shown by the Hi-C DNA contact mapping (Gibcus et al., 2018). When cells enter mitosis, condensin II forms large loops in prophase. Then, during prometaphase (after NEBD) condensin I forms smaller nested loops within a larger loop formed by condensin II. The formation of large loops causes steric repulsion between sister chromatids, which pushes them apart to facilitate their resolution (Goloborodko et al., 2016). The extended larger loops formed by condensin II would provide greater steric repulsion, thus facilitating sister chromatid resolution more efficiently (Figure 7, left). Given this, how could the precocious entry of condensin I into the nucleus in prophase interfere with the completion of sister chromatid resolution, which is dependent on condensin II? We propose two possible mechanisms for this as follows: First, if both condensin I and II form chromatin loops in prophase, both large loops and nested smaller loops inside of them would be formed at a similar timing. Thus, individual large loops would not be extended, compared to the normal situation. Then, some regions of sister chromatids might not push each other apart sufficiently due to reduced steric repulsion, leading to inefficient or incomplete sister chromatid resolution for some chromosomes (Figure 7, right). Second, if condensin I were driven into the nucleus in prophase, its chromatin binding might directly inhibit large loop formation by condensin II. For example, it is suggested that, after condensin II forms large loops along the spindle axis, condensin I sub-organises these loops into smaller loops (Gibcus et al., 2018; Liang et al., 2015; Walther et al., 2018), possibly through a mechanism involving Z-loop formation (where neighbouring condensin complexes bypass each other on the chromatin) (Elbatsh et al., 2019; Kim et al., 2020). However, if condensin II and condensin I work concurrently, the Z-loop formation may interfere with condensin II-dependent large loop formation along the spindle axis if they are structurally incompatible with each other.

**Figure 7:**
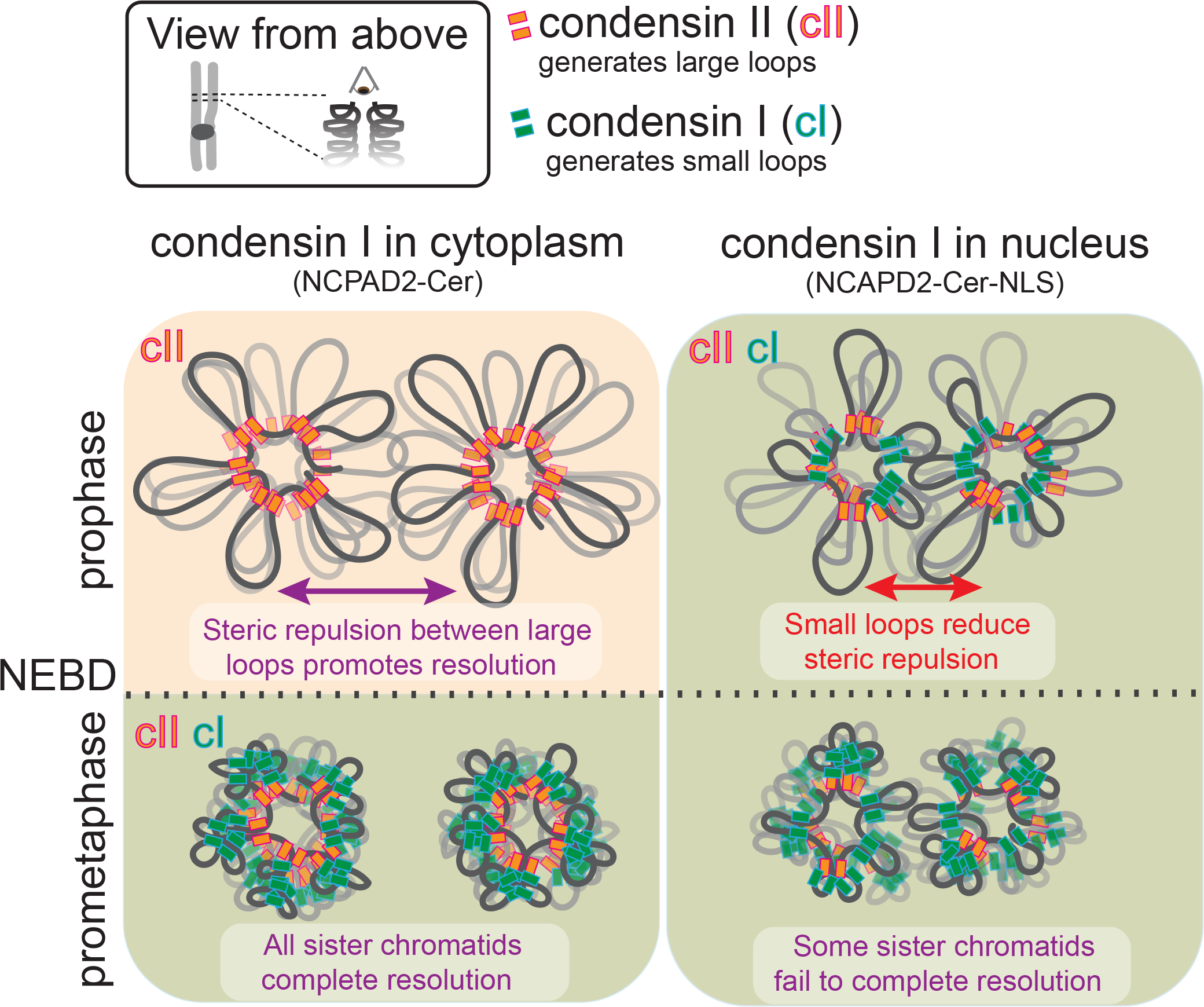
Model of how nuclear localization of condensin I affects mitotic chromosome reorganization. Here we illustrate the process of mitotic chromosome reorganization by viewing the cross-section of a pair of sister chromatids down their axes (illustrated in top left inset). Under normal circumstances (bottom left), only condensin II has access to chromosomes during prophase and large loops are formed, pushing sister chromatids apart due to steric repulsion (Goloborodko et al., 2016). At this stage, sister chromatid resolution is completed and sister chromosomes are shortened along their axes. Following nuclear envelope breakdown, small loops are formed by condensin I within the already formed large loops, making compacted (narrower) chromatids (Gibcus et al., 2018). In the situation where condensin I is present in the nucleus during prophase (bottom right), the premature formation of small loops, as well as large loops, will lead to reduced steric repulsion, resulting in incomplete sister chromatid resolution for some chromosomes. In this situation, at later stages of mitosis, whilst sister resolution is incomplete for some chromosomes, overall chromosome compaction looks relatively normal.

Although the majority of cells with NCAPD2-Cer-NLS proceeded in a timely manner to anaphase, some cells were arrested in metaphase, which was dependent on the SAC. What triggered this SAC-dependent metaphase arrest in these cells? We propose two non-mutually exclusive possible mechanisms: First, as sister chromatid resolution was often not completed with NCAPD2-Cer-NLS, the catenation checkpoint may have been engaged. It is known that the catenation checkpoint triggers metaphase arrest dependent on SAC regulators (Brownlow et al., 2014; Soliman et al., 2023). Second, condensin II at the centromere is important to maintain the centromere structure and its defect delays the onset of anaphase (Manning et al., 2010; Sacristan et al., 2024). It is possible that NCAPD2-Cer- NLS impaired the function of condensin II at the centromeres, leading to SAC-dependent metaphase arrest.

### Wider implications and human diseases

Several studies have demonstrated that disruption of either condensin I or condensin II has severe effects on chromosome segregation in anaphase (Green et al., 2012; Hudson et al., 2003; Ono et al., 2003; Umbreit et al., 2020). This is because both sister chromatid resolution by condensin II and chromosome compaction by condensin I are essential for chromosome segregation in mitosis. Here we show that the sequential action of first condensin II during prophase and then condensin I after NEBD is also critical to ensure proper chromosome segregation. Whilst in our experimental system we engineered translocation of condensin I to the nucleus such that it interacted with DNA precociously, there may be pathological situations leading to the same outcome.

One example might be nuclear envelope (NE) rupture in interphase or prophase, during which cytoplasmic proteins can enter the nucleus (Maciejowski and Hatch, 2020). NE rupture could also lead to entry of condensin I into the nucleus. NE rupture is usually repaired in a few minutes (De Vos et al., 2011; Vargas et al., 2012) whilst condensin I becomes active only in mitosis. So, if NE rupture occurs during interphase and its repair is followed by condensin I being pumped out of the nucleus, it might have no effects on chromosome organisation. However, if condensin I stays on chromosomes after the repair of the NE rupture in interphase, or if NE rupture happens during prophase, it may change chromosome reorganization in mitosis and may lead to chromosome missegregation and micronucleus formation. Because micronuclei provide less stable environments for chromosomes (Crasta et al., 2012; Zhang et al., 2015), a single rupture that allows condensin I to move into the nucleus precociously might have profound effects on genomic stability. NE rupture could also be associated with chromosome instability (CIN) as it is found in CIN+ cell lines such as HeLa and U2OS (Hatch and Hetzer, 2016; Vargas et al., 2012). In these cells, the entry of condensin I into the nucleus upon nuclear rupture might be part of an adverse reaction leading to chromosome instability.

Our results suggest different subcellular localization of condensin I and condensin II play important roles in their temporal coordination to ensure complete sister chromatid resolution and high-fidelity chromosome segregation. However, it is still unclear how their subcellular localization is regulated during the cell cycle. Meanwhile, various mutations in condensin I and condensin II subunits have been associated with cancer or other diseases (Baergen et al., 2019; Leiserson et al., 2015; Martin et al., 2016; Pang et al., 2022; Wang et al., 2021; Wang et al., 2018; Weyburne and Bosco, 2021). Many of these mutations may affect the intrinsic activity of the condensin complexes (i.e. independent of their subcellular activity) but others may affect their subcellular localization. It is important to elucidate how such mutations impair the functions of condensin complexes and how they lead to human diseases.

## Materials and Methods

### Cell culture

The human cell line HT-1080 (obtained from American Type Culture Collection) and derivative cell lines were cultured at 37°C and 5% CO2 under humidified conditions in DMEM (with L-glutamine), 10% FBS, 100 U/ml penicillin and 100µg/ml streptomycin that were all from Invitrogen. For live-cell microscopy, the above medium was replaced with Fluorobrite DMEM medium (Invitrogen) supplemented with 10% FBS, 2mM L-Glutamine, 1mM pyruvate and 25 mM Hepes.

Transfection of plasmids into HT-1080 derivative cell lines was carried out using Fugene HD (Promega) according to manufacturer guidelines. Briefly, cells were transfected in single wells of a 6-well dish using 3µl Fugene HD and 1µg plasmid (3:1 ratio). The selection agents G418 (Sigma; 300µg/ml) or zeocin (Invivogen; 200µg/ml) were introduced 48 hours after transfection.

Protein knockdown by siRNA was carried out using lipofectamine (Invitrogen) according to manufacturer guidelines. For cells grown in 2ml of medium in wells of a 6-well dish or in 3cm microscopy dishes 0.01nmol siRNA with 6µl lipofectamine and 200µl of Optimem (Invitrogen) were added. For cells grown in 300µl of medium in chambers of an 8-well microscopy slide (Ibidi) 1.5 pmol siRNA with 0.9µl lipofectamine and 30µl Optimem were added. The medium containing the siRNA was replaced along with fresh siRNA 24 hours later. Cells for analysis by western blotting or microscopy were taken between 48 and 60 hours after the first addition of siRNA. siRNAs were manufactured by Eurofins. The sequences were as follows; NCAPD2 (5’-CGUAAGAUGCUUGACAAUUTT-3’); control (scramble; 5’-UAACGACGCGACGACGUAATT-3’).

Cells were synchronized at the G1/S phase boundary by using a double-thymidine block. Briefly 0.1 to 0.2 x 10^6^ cells were seeded in 2 ml of medium in 6-well dishes or 3cm glass-bottomed microscope dishes (World Precision Instruments) or 0.04 x 10^6^ cells were seeded in 300µl of medium in chambers of an 8-well microscopy slide (Ibidi) 16 to 24 hours before treatment. Thymidine was added at a final concentration of 2.5mM and incubated for 16 hours. Thymidine was then removed and cells were washed with 3 x volume of fresh medium. Cells were incubated for a further 8 hours before 2.5mM thymidine was added.

Cells were then incubated for 12-16 hours and thymidine was then removed and cells were washes with 3 x volumes of fresh medium. After release cells were incubated for a further 8- 10 hours by which point they were reaching late G2/M phase.

Cells expressing CDK1as were synchronized at the G2/M boundary using 1NMPP1. Briefly 0.1 to 0.2 x 10^6^ cells were seeded in 2 ml of medium in 6-well dishes or 3cm glass-bottomed microscope dishes (World Precision Instruments) or 0.04 x 10^6^ cells were seeded in 300µl of medium in chambers of an 8-well microscopy slide (Ibidi) 16 to 24 hours before treatment.

1NMPP1 was added at a final concentration of 1µM and incubated for 12 to 16 hours. 1NMPP1 was then removed and cells were washed with 6 x volume of fresh medium to release cells into mitosis.

The Mps1/AuroraB inhibitor Reversine (Sigma; R3904) was used at a concentration of 100nM. The Mps1/TTK inhibitor CPD-5 (StressMarq Biosciences/2bScientific; SIH-183) was used at a concentration of 7.8nM. The DNA stain SiR-DNA (Spirochrome) was used at a concentration of 200nM and was added to cells approximately 10 hours before imaging.

### Plasmids

For stably expressing C-terminal Cerulean-tagged NCAPD2 in HT-1080 cells the plasmids pT3322 (NCAPD2-Cer-NLS) and pT3325 (NCAPD2-Cer) were generated. The important features of these plasmids are highlighted in Figure 1B and include C-terminal Cerulean tag with or without nuclear localisation signals, and also contains synonymous mutations of the NCAPD2 cDNA that inactivate an siRNA target site and a G418 resistance gene for selection in human cells. To create such plasmids, we first obtained a pcDNA3.1-based plasmid containing both the human NCAPD2 cDNA under the control of the constitutive CMV promoter and the G418 resistance gene for selection in vertebrate cells (Genscript; pcDNA3.1-NCAPD2).

To introduce the synonymous mutations at the siRNA site we cut out the original region from the pcDNA3.1-NCAPD2 plasmid by EcoRV/BmgBI restriction and replaced it with a similar DNA fragment (amplified using pcDNA3.1-NCAPD2 as template) containing the specific mutations. This DNA fragment was generated by PCR amplification of the 5’ region containing the EcoRV site and the 3’ region including BmgBI using primer pairs 6374 (5’- AGA ACT TCT TCA ATG AGC TCT CC -3’) / 6375 (5’- A**G**T T**A**T C**C**A **A**CA T**T**T T**C**C **T**GA GGC CTC GCT CTG TGA GGG GC -3’) and 6376 (5’- **A**G**G** AA**A** ATG **T**T**G** GA**T** AA**C** TTT GAC TGT TTT GGA GAC AAA CTG -3’) / 6377 (5’- AAT CTC AAG CTC CTT GAT TCC AT -3)’, respectively. Note that the primers 6375 and 6376 were designed with the synonomous mutations incompatible with the siRNA sequence (mutations indicated in bold). These two products were fused in the second round of PCR (using primer pair 6374 / 6377) and this product was then cloned into the original pcDNA3.1-NCAPD2 plasmid to generate the plasmid designated pT3310.

To introduce the DNA encoding the cerulean tag with or without nuclear localisation signals we cut out a region of pT3310 (including the 5’ coding sequence of NCAPD2 and the 3’ non-coding flanking sequence) by BmgBI/DraIII restriction and replaced it with either of two PCR products in which a cerulean tag with or without a NLS had been introduced in-frame with the NCAPD2 coding sequence and immediately upstream of the 3’ flanking region. The 5’ coding and 3’ non-coding flanking regions were amplified using template DNA provided by pcDNA3.1-NCAPD2. The 5’ coding region was generated using primer pair 6378 (5’- GTT TTG GAG ACA AAC TGT CAG ATG -3’) / 6379 (5’- AGC AGC TGA AGC GGC TGA GGC TCC GGA TCT GTG CCT GCG AGC CG -3’), the 3’ non-coding flanking region (no NLS) using primer pair 6383 (5’- GAC CCT AAG AAA AAA CGT AAA GTG GAT CCG AAG AAA AAG AGG AAA GTC TGA TAA ACC CGC TGA TCA GCC TCG -3’) / 6385 (5’- TCC ACT ATT AAA GAA CGT GGA CTC -3’) and the 3’ flanking region with NLS using primer pair 6384 (5’- GCG TAT CGA TCC TGA TAA ACC CGC TGA TCA GCC TCG -3’) / 6385 (as above). The fragment encoding the Cerulean gene containing 3xNLS was amplified from the template pCerulean-PCNA-19-SV40NLS-4 (a gift from Michael Davidson; Addgene plasmid # 55437) using primer pair 6380 (5’- GGA GCC TCA GCC GCT TCA GCT GCT CCG GTC GCC ACC ATG GTG AG -3’) / 6381 (5’- CAC TTT ACG TTT TTT CTT AGG GTC TAC CTT TCT CTT CTT TTT TGG ATC GGA TCG ATA CGC GTA GCC GC -3’) and a version that excluded the NLS region was amplified using primer pair 6380 (as above) / 6382 (5’- GCG GGT TTA TCA GGA TCG ATA CGC GTA GCC GC -3’). These PCR products were fused together with the 5’ coding and 3’ non-coding fragments in a further two rounds of PCR and then cloned into pT3310 to generate plasmids designated pT3314 (containing NCAPD2-Cer- NLS) or pT3316 (containing NCAPD2-Cer). The DNA region containing NCAPD2-Cer-NLS or NCAPD2-Cer genes as well as the G418 resistance gene were subcloned, by restriction with BmgBI and DraIII, into the plasmid pMK243 (Natsume et al., 2016) replacing both its TIR1 expression cassette and existing resistance gene found between AAVS1 homology regions, to generate plasmids designated pT3322 and pT3325 respectively. These two plasmids were used for transfection and generation of stable human cells expressing NCAPD2-Cer or NCAPD2-Cer-NLS.

### Cell lines

The human HT-1080–derived cell line containing *tet* operator and *lac* operator arrays, separated by 250 kbp of DNA and expressing TetR-4mCherry and EGFP-LacI (designated TT75) was described previously (Eykelenboom et al., 2019). Derivatives of TT75 were generated that expressed NCAPD2-Cer or NCAPD2-Cer-NLS by transfection with plasmids pT3325 and pT3322, respectively, and selection with 300µg/ml G418. These cell lines were designated TT204 and TT234 respectively. Alternative NCAPD2-Cer or NCAPD2-Cer-NLS stable cell lines, whose origins are the clones (cell colonies after transfection) different from TT204 and TT234, were also used in some experiments and were designated TT168 and TT170, respectively. A derivative of the TT75 cell line was generated, whose endogenous CDK1 genes were disrupted and that expressed *Xenopus* CDK1as, using plasmids pX330_human CDK1, pCDK1as_T2A_Zeo (gifts from William Earnshaw; Addgene 118597 and 118596, respectively) and pCMV(CAT)T7-SB100 (a gift from Zsuzsanna Izsvak; Addgene 34879) (Hochegger et al., 2007; Mates et al., 2009; Saldivar et al., 2018). The cell line was generated according to published methodology using 200µg/ml zeocin selection (Saldivar et al., 2018). This cell line was designated TT206.

### SDS PAGE and western blotting

For western analysis, whole cell extracts were prepared from cells grown in 6-well dishes and lysed in 30–50 μl of lysis buffer (20 mM Hepes, pH 7.6; 400 mM NaCl; 1 mM EDTA; 25% glycerol; 0.1% NP-40) containing protease inhibitors (cOmplete EDTA- free; Roche). Bradford reagent (Thermo Fisher Scientific; 1863028) was used to measure protein concentration of each lysate and 50 μg of total protein of each was run on precast Bis-Tris 4–12% gradient gels (Invitrogen) and protein subsequently transferred to polyvinylidene fluoride membrane (Amersham). Membranes were first blocked in PBS containing 2% BSA and then were incubated overnight at 4°C with primary antibodies diluted in PBS containing 2% BSA and 0.05% (w/v) sodium azide. Membranes were then washed 3 times with PBS containing 0.1% Tween20 (Sigma) and then were incubated for 2 hours at room temperature with flouorescently-labelled secondary antibodies diluted in PBS containing 1% milk powder. Membranes were then washed 3 times with PBS containing 0.1% Tween20 (Sigma) and signal from the secondary antibody was detected using a LI-COR Odyssey CLx. Primary antibodies were used as follows: NCAPD2 (Sigma; HPA036947), 1 in 5,000; actin (Sigma; A5441), 1 in 20,000. Secondary antibodies were all used 1:10,000 and were: donkey-anti-mouse-800CW (LI-COR; 926-32212); donkey-anti-rabbit-680RD (LI-COR; 926–68073).

### Live-cell microscopy and image analysis

Time-lapse images were collected at 37°C with 5% CO2 using a DeltaVision ELITE microscope (Applied Precision). We used an apochromatic 100x objective lens (Olympus; numerical aperture: 1.40) or 60x objective lens (Olympus; numerical aperture: 1.42) to minimize longitudinal chromatic aberration. We also routinely checked lateral and longitudinal chromatic aberration using 100-nm multi-color beads. We did not detect any chromatic aberration between the colors observed in the current study. For signal detection we used a sCMOS camera (PCO Edge).

For tracking *lac* and *tet* operator arrays in live cells, or imaging fixed cells we used a 100x objective lens and acquired 25 z-sections 0.5 μm apart with 2x2 binning. For live cells Cerulean, EGFP, and mCherry signals were discriminated using the dichroic CFP/YFP/mCherry (52-850470-000 from API) or EGFP and mCherry signals were discriminated using the dichroic DAPI/FITC/mCherry/Cy5 (52-852112-001 from API). For fixed cells, we used a 100x objective lens and acquired 32 z-sections 0.25µm apart with 2x2 binning. DAPI, Alexa488, Alexa565 and Alexa647 were discriminated using the dichroic DAPI/FITC/TRITC/Cy5 (52-852111-001 from API). For tracking stained chromosomes in live cells we used a 60x objective lens and acquired 12 z-sections 1.5µm apart with 4x4 binning. SiR-DNA (Spirochrome; emission 674 nm) and EGFP signals were discriminated using the dichroic DAPI/FITC/mCherry/Cy5 (52-852112-001 from API).

After acquisition, images were deconvolved using softWoRx software with enhanced ratio and 10 iterations. Analysis of individual cells was performed using Imaris software (Bitplane). Statistical analyses were performed using Graphpad Prism 6.0 software.

Imaris was used to automatically assign xyz coordinates to the center of mass of the fluorescent dots (the centroid), associated with each *lac* or *tet* operator array (LacO and TetO dots). From these coordinates, distances between LacO and TetO dots (Figure S2A; distances a, b, c, d, r and g) and, where four distinct dots were observed, the angle (LL-TT angle) between the vector connecting two LacO dots (vector LL) and the vector connecting two TetO dots (vector TT) were calculated. Because the order of dots in each vector is undefined, the angle was limited to be between 0° and 90°. Specifically, the angle was calculated as acos|*μ*| where *μ* is the cosine angle between vectors **LL** and **TT**, *μ* = **LL** ·**TT**/(‖**LL**‖‖**LL**‖). We then set definitions of the configurations of LacOs and TetOs (non-resolved, partially resolved, resolved or compacted state: blue, brown, pink or red pattern respectively; Figure 2B) in a similar manner to our previous publication (Eykelenboom et al., 2019), using a purpose-built R Shiny app, which is available at https://github.com/bartongroup/MG_ChromCom2. The four configurations were defined as follows: For both blue “non- resolved” and brown “partially resolved” states, either LacO or TetO was observed as a single fluorescent object or 2 objects whose centers were separated <0.2 µm. To distinguish blue “non-resolved” and brown “partially resolved” states, we looked at the second operator dot and (a) if it was observed as a single fluorescent object or appeared as two fluorescent objects whose centers were separated by <0.75 μm, we classified it as the blue “non-resolved” state, or (b) if it was observed as two fluorescent objects whose centers were separated by >0.75 μm, we classified it as the brown “partially resolved” state. For both pink “resolved” and red “compacted” states, both LacO and TetO appeared as two separate fluorescent objects (four objects in total; two LacOs and two TetOs). First, we assumed the shorter distance between a+b and c+d defined which LacO and TetO were on the same chromatid (Figure S2A) and only considered these two distances for our assignment. To distinguish pink “resolved” and red “compacted” states, we first looked at two LacO-TetO distances, and (i) if both LacO-TetO distances were >0.4µm apart, we classified them as the pink “partially resolved” state, or (ii) if both LacO-TetO distances were <0.4µm apart, we classified them as the red “compacted” state. If neither (i) nor (ii) applied, we classed them as borderline cases. Because we had noticed that in earlier stages during resolution (prophase), when LacO-TetO distances were larger, the LL-TT angle tended to be larger and closer to 90°, we used this angle to discriminate the pink “resolved” and red “compacted” patterns in borderline cases, as follows: if either of the LacO-TetO distances was >0.4µm and the LL-TT angle was >45°, we classified it as the pink “resolved” state. In all other cases we classified them as the red “compacted” state.

### Fixed-cell microscopy and image analysis

For fixed-cell immunofluorescence cells were grown and treated in 3cm fluorodishes (WPI) and fixed using acetone-methanol (Hirota et al., 2004) or by paraformaldehyde. For acetone-methanol fixation cells were washed once with 2ml PBS and then replaced with 2ml of ice-cold acetone-methanol (1:1) for 2 minutes followed by 2 minutes each with 2ml room temperature PBS containing decreasing amounts of methanol [95%, 90%, 80%, 70% and 50% (v/v)]. For paraformaldehyde fixation cells were washed once with 2ml PBS which was replaced with 2ml of room temperature 4% PFA in PBS (pH6.8) for 10 minutes. The cells were then washed with 2ml PBS three times. Paraformaldehyde fixed cells were permeabilized by treatment with 2ml of room temperature PBS containing 0.5% Triton for 10 minutes. The cells were then washed with 2ml PBS twice. After fixation (and permeabilization) and before immunostaining cells were incubated with PBS containing 2- 3% BSA at 4°C for at least 2 hours. Antibodies were then diluted appropriately in PBS containing 2-3% BSA and then incubated with fixed cells overnight at 4°C. The cells were then washed with 2ml PBS three times before incubation with fluorescently-labelled secondary antibodies diluted appropriately in PBS containing 2-3% BSA overnight in the dark at 4°C. Before imaging the cells were then washed with 2ml PBS three times. In some cases these cells were then imaged directly. In cases where DNA visualisation was required the cells were mounted with coverslips and 25µl Prolong Gold Antifade containing DAPI (Invitrogen) which was left to polymerise in the dark at room temperature for at least 2 hours before imaging. Primary antibodies were used as follows NCAPG (Santa Cruz; sc-515297) 1:200, NCAPH (Novus Bio; NBP1-88345) 1:200, Alpha-Tubulin (Merck; mab1864) 1:1,000; ACA (Antibodies incorporated; 15235) 1:400. Secondary antibodies were all used at 1:1,000 and were Donkey anti-human IgG-Alexa647 (Jackson ImmunoResearch 709-605-149), Donkey anti-rat Alexa488 (Invitrogen; A21208), Donkey anti-rabbit Alexa647 (Abcam; ab150075), Goat anti-mouse Alexa647 (Abcam; ab150119).

## Acknowledgements

We thank members of Tanaka lab for discussion and G. Barton for supervising the Data Analysis Group. We also thank M. Kanemaki lab for plasmid pMK243; M. Davidson lab for plasmid pCerulean-PCNA-19-SV40NLS-4; W. Earnshaw lab for plasmids pX330_human CDK1 and CDK1as_T2A_Zeo and Z. Izsvak lab for plasmid pCMV(CAT)T7-SB100. This work was supported by BBSRC project grant (Ref BB/S007768/1), MRC project grant (Ref MR/T046880/1) and Wellcome Trust Investigator Award (Ref 219418/Z/19/Z).

The authors declare no competing financial interests.

## Author contributions

Conceptualization: T.U. Tanaka, J.K. Eykelenboom; Methodology: J.K. Eykelenboom, M. Gierliński; Investigation; J.K. Eykelenboom, M. Gierliński, Z. Yue; Formal analysis: J.K. Eykelenboom, M. Gierliński; Software: M. Gierliński; Visualization: J.K. Eykelenboom, M. Gierliński, T.U. Tanaka; Writing (original draft): J.K. Eykelenboom; Writing (review and editing): T.U. Tanaka, J.K. Eykelenboom; Project administration, supervision and funding acquisition: T.U. Tanaka.

---

**Figure S1:**
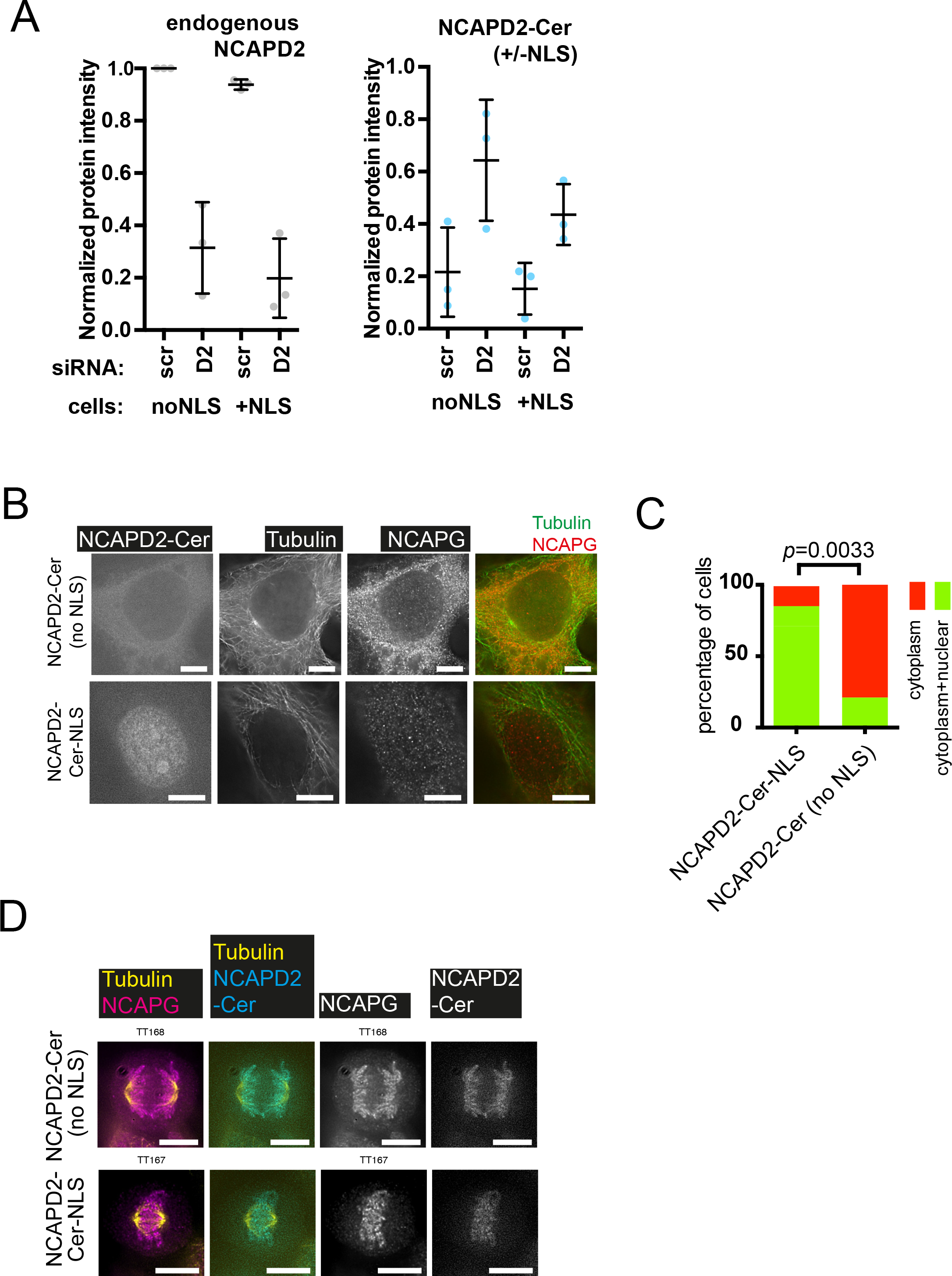
Characterisation of cell lines expressing NCAPD2-Cer or NCAPD2-Cer-NLS. *(Related to* Figure 1). **(A)** Quantification of western blotting for endogenous NCAPD2 (left hand panel) and NCPAD2-Cer (+/-NLS) (right hand panel) following treatment with non-specific (scramble; scr) or NCAPD2 (D2) siRNA. Graphs show the mean protein signal intensities, normalized to endogenous NCAPD2 in control cells treated with non-specific siRNA. Bars indicate the standard deviation from three independent experiments. **(B)** Localization of condensin I subunits in fixed interphase cells stably expressing either NCAPD2-Cer (top) or NCAPD2-Cer-NLS (bottom) after treatment with NCAPD2 siRNA for 48 hours. NCAPD2-Cerulean was visualised directly whilst NCAPG and Tubulin were visualised with immunofluorescence staining. The top and bottom panels show typical interphase cells displaying cytoplasmic or nuclear/cytoplasmic localisation of NCAPG, respectively. Scale bar 10 µm. **(C)** The graph shows quantification of the patterns from (B) for the indicated stable cell lines. The *p* value was obtained using a chi-square test. n = 6 and 9 for each cell line respectively. **(D)** Fixed mitotic cells expressing NCAPD2-Cer (top) or NCAPD2-Cer-NLS (bottom). NCAPD2-Cer, Tubulin and NCAPG were visualized as in Figure 1F, on mitotic chromosomes. Scale bar 10µm.

**Figure S2:**
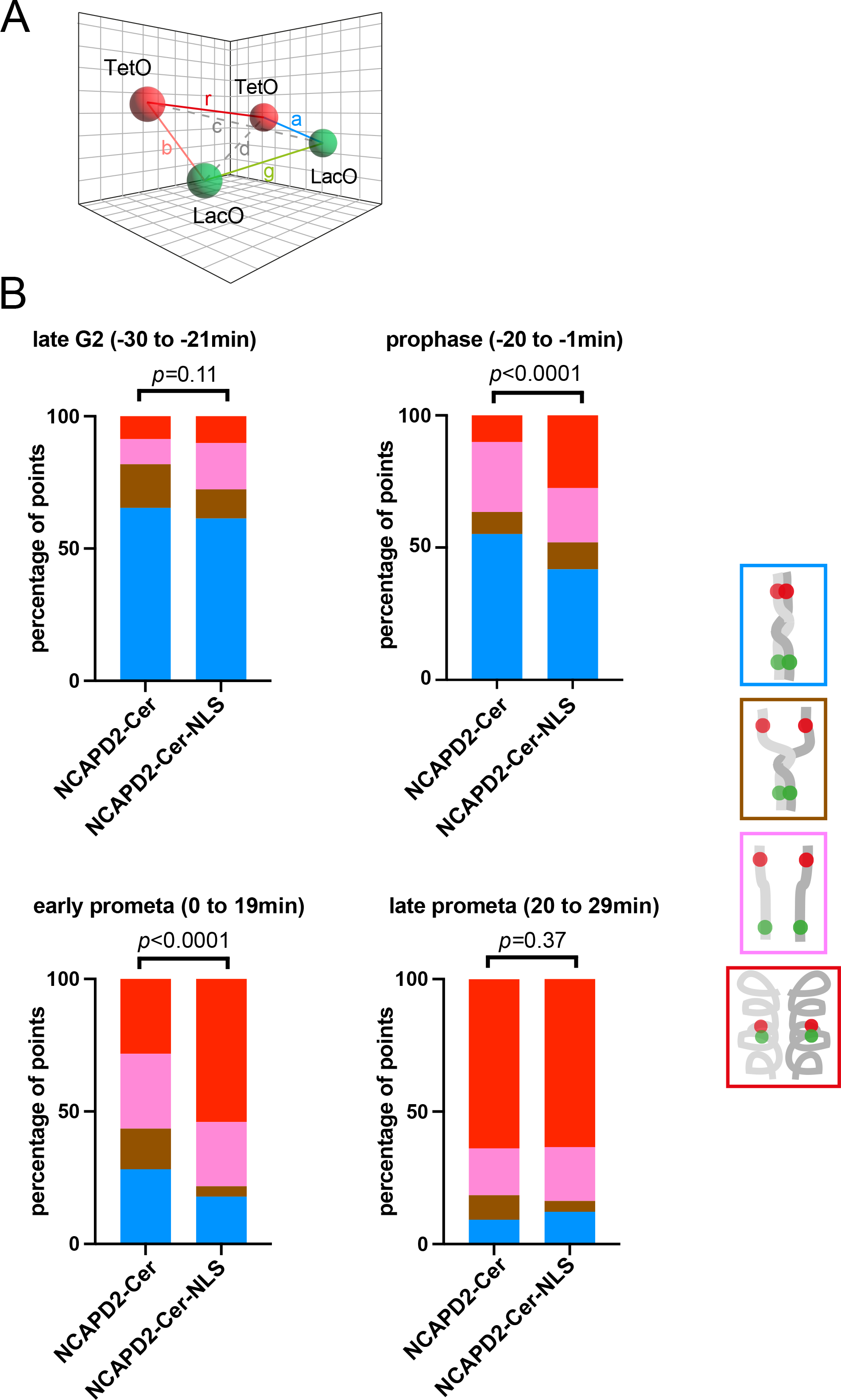
Analyses of *tet*- and *lac*-operator arrays to determine mitotic chromosome behaviours. *(Related to* Figure 2). **(A)** Relevant distances between the TetR-mCherry and LacI-GFP dots as calculated from xyz coordinates. Note that the full 3-dimensional relationship was considered when making measurement of each distance. For detailed explanation of distinguishing different states see Materials and Methods. **(B)** Graph shows the proportional representation of each state during late G2 phase (-30 to -21 mins), prophase (-20 to -1 mins), early prometaphase (0 to 19 mins) and late prometaphase (20 to 29 mins) for NCAPD2-Cer or NCAPD2-Cer-NLS cells. The data was taken from Figure 2C and D. The prophase data is a copy of that shown in Figure 2F. The *p* values were obtained using chi-square tests. For late G2 phase n = 228 and 127, prophase n = 419 and 515, for early prometaphase n = 363 and 177 and for late prometaphase n = 123 and 119, in each case for NCAPD2-Cer and NCAPD2- Cer-NLS cells respectively.

**Figure S3:**
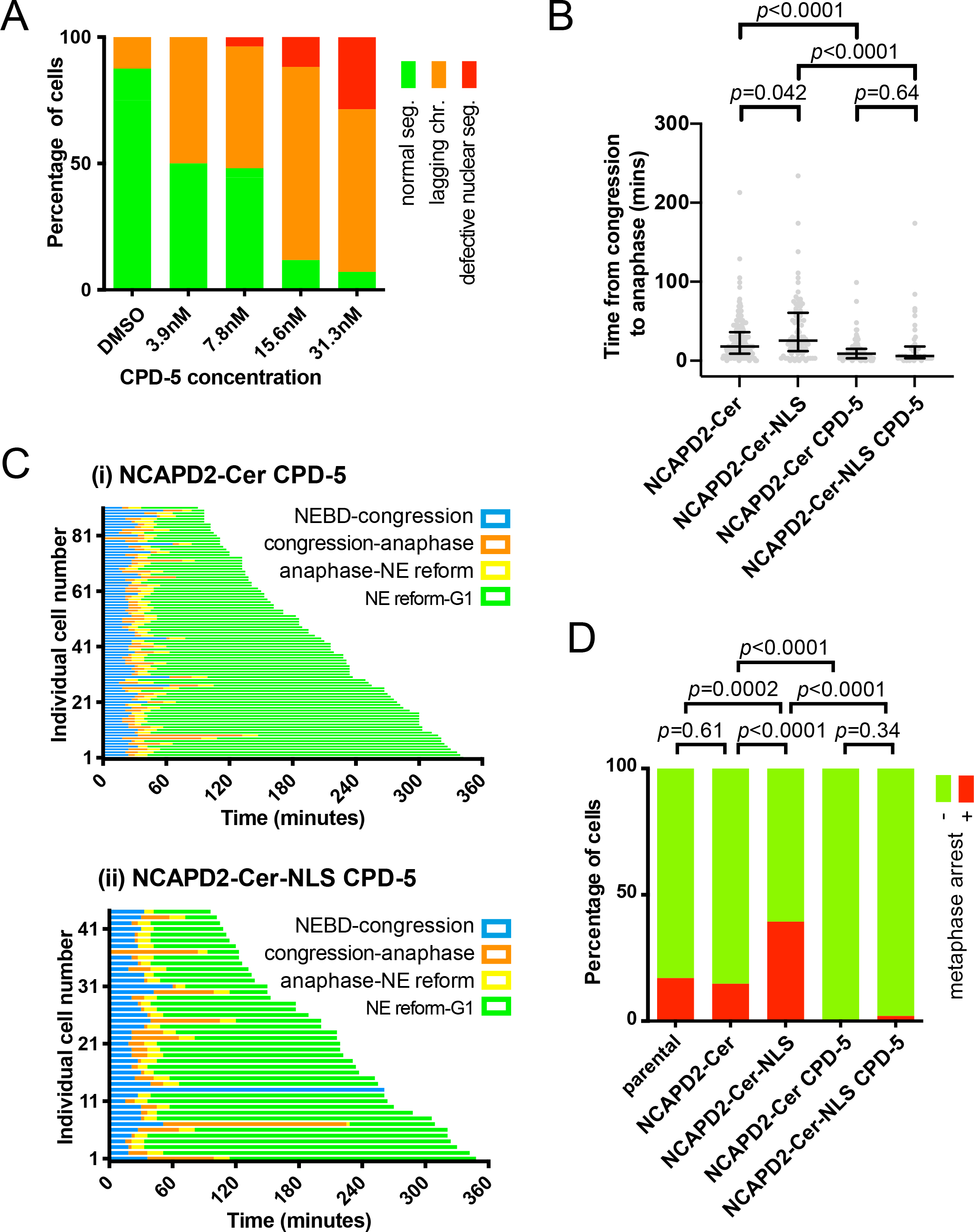
Analysis of mitotic progression of NCAPD2-Cer and NCPAD2-Cer-NLS cells in the presence of CPD-5. *(Related to* Figure 4)**. (A)** Analyses of chromosome segregation in the parental cells expressing *Xenopus* CDK1as (see Materials and Methods) that were synchronized at the G2-M phase boundary with 1NMPP1. After 1NMPP1 washout cells were treated with different concentrations of CPD-5 as indicated, microscopy images were taken every minute, and anaphase was observed. NEBD and the nuclear envelope (NE) reform were identified as in Figure 4A. Chromosomes were visualized using SiRDNA. The outcomes of anaphase quantified in this experiment are shown in Figure 5A and B. **(B)** Time elapsed between completion of congression of chromosomes on the metaphase plate and the onset of anaphase of individual cells treated or not with 7.8nM CPD-5. Timing of the two events was measured by visualizing chromosomes as in Figure 4A. Horizontal lines and bars indicate the median and interquartile range for each condition. The *p* values were obtained using Mann-Whitney U tests. n = 165, 82, 91 and 43 for each condition from left to right. **(C)** Mitotic progress of individual NCAPD2-Cer or NCAPD2-Cer-NLS cells treated with siRNA against endogenous NCAPD2 and 7.8nM CPD-5 (*y* axis) plotted against time (*x* axis) and aligned relative to NEBD (defined as time zero). The colored lines, that use the same color codes as Figure 4A, refer to trajectories of individual cells through mitosis. The end of the line indicates the end of the time-lapse for the cell. The number of analyzed cells were 91 and 44 for NCAPD2-Cer and NCAPD2-Cer-NLS cells, respectively. **(D)** Quantification of the number of NCAPD2-Cer, NCAPD2-Cer-NLS or parental cells (i.e, with no NCAPD2-Cer construct) that did not progress further than metaphase during the time-lapse acquisition. The graphs are based on the data in Figures 4B and S4B. The left 3 bars are the same graphs as Figure 4C. The *p* values were obtained using Fisher’s exact test.

**Figure S4:**
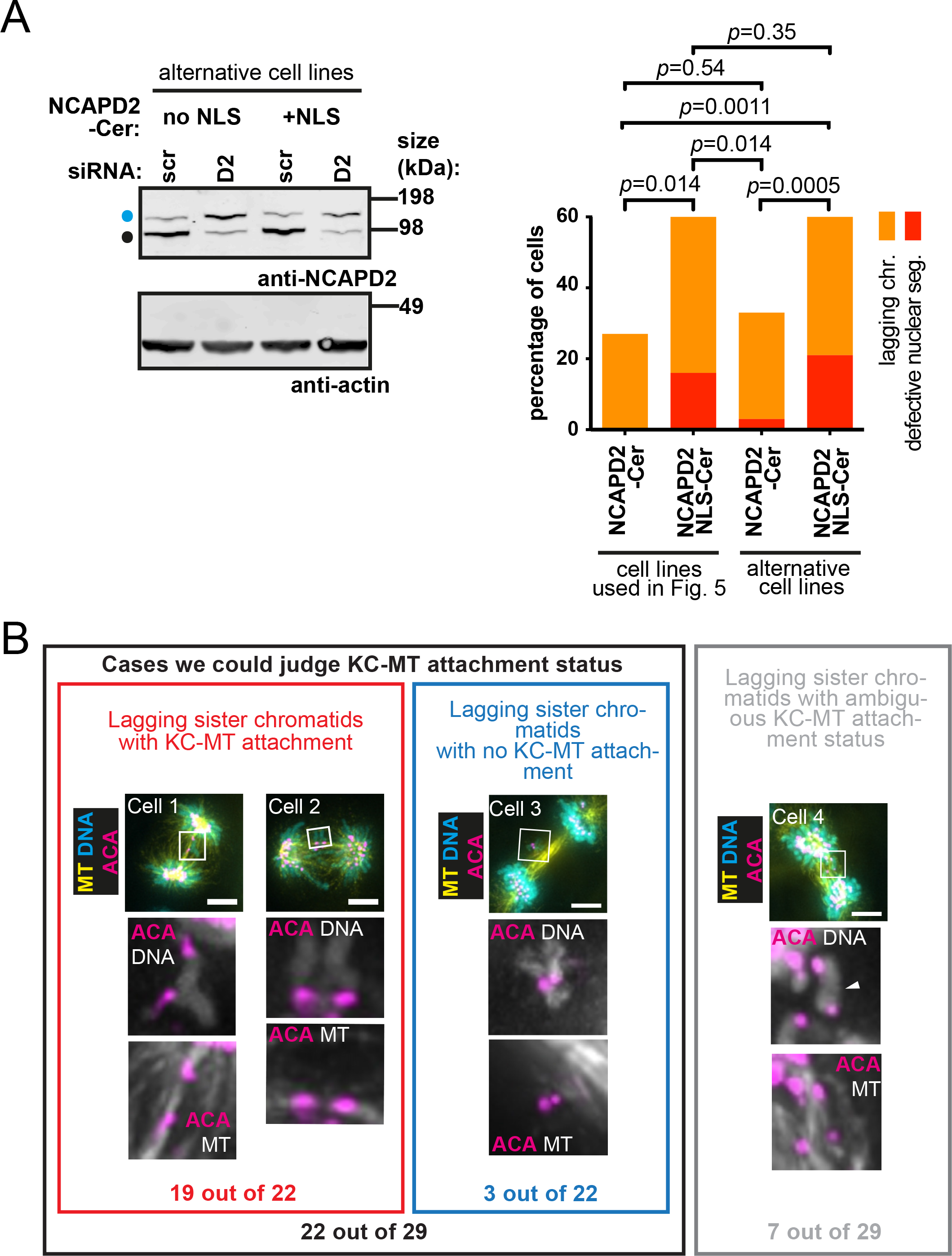
Further analyses of chromosome segregation during anaphase in NCAPD2-Cer-NLS cells. *(Related to* Figure 5 *and 6)* **(A)** Analysis of chromosome segregation during anaphase in alternate cell lines expressing NCAPD2-Cer or NCAPD2- Cer-NLS. Left; a western blot with an anti-NCAPD2 antibody for proteins from alternative NCAPD2-Cer and NCAPD2-Cer-NLS cell lines after treatment with NCAPD2 siRNA for 48 hours. The endogenous protein (NCAPD2; sensitive to the siRNA) is indicated by a small black circle whilst NCAPD2 tagged with Cerulean (NCAPD2-Cer; insensitive to the siRNA) is indicated by a blue circle. Actin is shown as a loading control. Right; quantification of chromosome segregation defects for two independent NCAPD2-Cer and two independent NCAPD2-Cer-NLS cell lines. Cells were treated, imaged and categorized according to Figure 5A and B. In each cell line. The cells that performed anaphase without segregation defects make up the remainder percentages but are not shown on the graph. The *p* values were obtained using chi-square test for trends. The number of analyzed cells, from left to right, were 15, 25, 30 and 33. **(B)** Analysis of lagging chromosomes with centromere pairs in anaphase of NCAPD2-Cer-NLS cells. Fixed-cell analysis of centromeres and chromosomes during anaphase. Cells were prepared for immunofluorescence staining as described in Figure 6A. Lagging chromosomes with paired centromeres were identified in NCAPD2-Cer- NLS cells. Cell 1 and 2 displayed typical examples of pairs of lagging sister chromatids with microtubule attachments from opposite spindle poles. Cell 3 contained a pair of lagging sister chromatids that were not attached to microtubules from opposite spindle poles. Cell 4 contained a pair of lagging sister chromatids (indicated by white arrowhead) whose kinetochore-microtubule (KC-MT) attachment status was unclear. Scale bars 5µm. Cell 1 is the same cell as shown in Figure 6A (cell number 5). Cell 3 is the same cell as shown in Figure 6A (cell number 4). We analyzed 29 pairs of lagging sister chromatids (with immunostaining of both ACA and microtubule), from the data set analysed in Figure 6.

